# Increased Nuclear Susceptibility to Mechanical Stress in Phospholamban R14del Cardiomyopathy: A New Perspective on Disease Heterogeneity

**DOI:** 10.64898/2026.10.23.761407

**Authors:** Jort S. A. van der Geest, Atze van der Pol, Leon Hermans, Achmed Al-khaqani, Jeroen van Asten, Marlinne A.M.E. van Roessel, Willem B. van Ham, Anneline S.J.M. te Riele, Alain van Mil, Vasco Sampaio-Pinto, Pieter A. Doevendans, Zhiyong Lei, Vito Conte, Joost P. G. Sluijter, Carlijn V.C. Bouten

**Affiliations:** Department of Cardiology and Experimental Cardiology Laboratory, Division of Heart & Lungs, University Medical Centre Utrecht, Utrecht, the Netherlands; Regenerative Medicine Centre Utrecht, Circulatory Health Research Center, University Utrecht, University Medical Centre Utrecht, Utrecht, the Netherlands; Department of Biomedical Engineering, Eindhoven University of Technology, Eindhoven, The Netherlands; Institute for Complex Molecular Systems, Eindhoven University of Technology, Eindhoven, The Netherlands; Department of Central Diagnostic Laboratory-Research, University Medical Center Utrecht, Utrecht University, Utrecht, the Netherlands; Department of Medical Physiology, Division of Heart & Lungs, University Medical Centre Utrecht, Utrecht, the Netherlands; Netherlands Heart Institute (NLHI), Utrecht, the Netherlands; Central Military Hospital (CMH), Utrecht, The Netherlands

**Keywords:** PLN R14del cardiomyopathy, Nuclear stress, Cell mechanical stimulation, Nuclear abnormalities

## Abstract

**Background:** Clinical presentation of phospholamban (PLN) R14del cardiomyopathy is highly variable, from asymptomatic carriers to rapidly progressing heart failure. which cannot be fully explained by its known disturbance of sarcoplasmic reticulum calcium handling. Nuclear calcium signalling, shaped by sarcoplasmic reticulum calcium release, plays a key role in regulating gene expression and the cellular response to mechanical stress. However, how the R14del variant affects nuclear signalling under mechanical loading remains unclear.

**Objective:** To uncover the impact of the PLN R14del variation on mechanosensing, nuclear calcium handling and its link to the variable clinical representation of this cardiomyopathy.

**Methods:** Human induced pluripotent stem cell-derived cardiomyocytes (hiPSC-CMs) carrying the PLN R14del variant, an isogenic control, and wildtype control were subjected to mechanical stimulation by cyclic stretch or cultured on substrates with physiological (10-20 kPa) and pathological (65-85 kPa) stiffness. Nuclear morphology, cytoskeletal organisation and nuclear body formation were assessed via immunofluorescence. Traction force microscopy was used to assess mechanical stress and force generation. Cardiac tissues from PLN R14del cardiomyopathy patients, dilated cardiomyopathy patients, and healthy controls were analysed under mechanical loading (at left ventricular assist device implantation) and unloading (at heart transplantation). Living myocardial slices from a PLN R14del patient were maintained under overloading or physiological loading conditions.

**Results:** PLN R14del hiPSC-CMs showed impaired nuclear calcium handling and, under mechanical stress, exhibited impaired mechanotransduction and elevated nuclear bodies. A strain-avoidance response was triggered but failed to prevent apoptosis. Nuclear abnormalities were confirmed in patient myocardial slices and cardiac biopsies which normalized upon relative mechanical unloading.

**Conclusion:** PLN R14del increases nuclear susceptibility to mechanical stress, linking nuclear dysfunction to new disease insights and highlighting potential for mechanobiology-based therapeutic interventions.

**Condensed abstract:** PLN R14del cardiomyopathy presents with highly variable clinical severity, which cannot be fully explained by its known disturbance of sarcoplasmic reticulum calcium handling. We investigated whether the R14del variant impairs nuclear calcium signalling and mechanosensing, thereby contributing to disease heterogeneity. Using PLN R14del, isogenic control, and wildtype hiPSC-CMs subjected to cyclic stretch and substrates of physiological and pathological stiffness, we demonstrate that PLN R14del cardiomyocytes exhibit impaired nuclear calcium handling, elevated nuclear damage, disrupted cytoskeletal organisation, and failed mechanotransduction under mechanical stress. Although a strain-avoidance response was triggered, it was insufficient to prevent apoptosis. These findings were confirmed in living myocardial slices and cardiac biopsies from PLN R14del patients, where nuclear abnormalities were most pronounced under mechanical loading and reversed upon unloading. Together, these data establish increased nuclear susceptibility to mechanical stress as a novel pathological feature of PLN R14del cardiomyopathy, offering new insight into disease heterogeneity and highlighting mechanobiology-based therapeutic opportunities.

## Introduction

Genotype-informed care is now central to cardiomyopathy management, with current guidelines advocating pathways in which genetic testing directly informs diagnosis, risk stratification, and family management^1, 2^. Within this framework, phospholamban (PLN)-cardiomyopathy caused by deletion of arginine at position 14 (c.40_42delAGA, p.Arg14del, hereafter referred to as R14del) variant is recognized as a distinct disease entity with dedicated diagnostic and risk-prediction approaches^3, 4^. The R14del founder variant was initially identified in the Netherlands and in Greece^5, 6^. From these regions, the variant has spread around the globe, having been reported in Belgium^7^, Canada^8^, China^9, 10^, Germany^11^, Japan^12^, Norway^7^, Spain^13^, the United Kingdom^7^, and the United States^14^, underscoring its clinical importance.

PLN regulates cardiac contraction by inhibiting sarcoplasmic reticulum Calcium-ATPase (SERCA) in its unphosphorylated state, thus limiting calcium reuptake during diastole, and thereby delaying cardiac relaxation.^15^ This inhibition is relieved upon phosphorylation by protein kinase A (PKA). Importantly, the R14del genetic variant disrupts this regulatory domain, impairing PLN phosphorylation and thereby altering its interaction with SERCA.^16^ Despite extensive research, the precise mechanism of the PLN R14del variant on calcium handling remains controversial, with reports ranging from SERCA super-inhibition to a loss of inhibitory function.^6, 17–23^ More recent work points beyond the effect on calcium kinetics to structural pathology of the sarcoplasmic reticulum (SR) with aberrant SR microdomain organisation^23, 24^. This supports the idea that the pathogenic effects of R14del extend beyond SR calcium cycling alone, highlighting the need to explore additional mechanisms that may contribute to the disease phenotype.

Notably, the PLN R14del variant is associated with a characteristic myocardial remodelling, including fibrosis and fibrofatty replacement^25–27^, together with cellular stress pathways including impaired autophagy resulting in perinuclear aggregates^23, 28^, mitochondrial dysfunction^18, 22^, and activation of the unfolded protein response (UPR).^29, 30^ Clinically, PLN-cardiomyopathy exhibits incomplete, age-dependent penetrance, with many carriers remaining asymptomatic^5, 31^. Once disease manifests, progression can be rapid, with a high burden of malignant ventricular arrhythmias, and progressive left-ventricular dysfunction that may evolve to biventricular failure and sudden cardiac death^5, 31, 32^. This spectrum, from lifelong asymptomatic carriage to rapid deterioration, illustrates the currently unexplained heterogeneity in disease severity and progression.^5, 32^

While the canonical role of PLN in SR calcium handling is well established, recent studies have hinted at non-canonical functions of PLN, particularly involving the nuclear envelope. PLN appears to exhibit unique trafficking patterns that differ from other SR proteins and interacts with nuclear-specific components, suggesting potential involvement in nuclear signalling pathways.^33, 34^ These observations invite investigation into how PLN R14del might alter nuclear behaviour, especially under mechanical stress.

The nucleus is increasingly recognized as a key mediator of mechanotransduction, the process by which cells sense and convert mechanical forces into biochemical signals and adaptive responses.^35^ This is particularly relevant in the heart, which is continuously subjected to contractile strain and hemodynamic loading. Importantly, calcium is a central messenger in mechanotransduction signalling, regulating not only contraction, but also gene expression, nuclear remodelling, and chromatin dynamics.^36^ Nuclear calcium handling facilitates nuclear softening and remodelling under mechanical strain, protecting it against stretch-induced damage.^37^

Given the mechanosensitive nature of the cardiomyocyte nucleus and the central role of calcium in this process, we hypothesize that PLN R14del impairs nuclear calcium dynamics and thereby nuclear mechanotransduction. As a consequence, this disruption can contribute to altered nuclear morphology, impaired stress adaptation, and downstream gene dysregulation. This may be especially detrimental in the fibrotic heart of PLN R14del patients, where matrix stiffening amplifies mechanical loading on the cardiomyocyte nucleus.^25–27^ We further speculate that differential mechanosensitivity among PLN R14del patient cells contributes to the variable disease presentation seen in patients, ranging from severe biventricular failure to asymptomatic carrier states.

In this study, we investigated the impact of the PLN R14del variant on the responsiveness of cardiomyocytes to mechanical stress, focusing on calcium signalling, nuclear morphology, cytoskeletal adaptations, and nuclear stresses. To this end, we employed a comprehensive set of human-relevant models, each mimicking a different aspect of mechanical loading. Mechanical loading was here defined as conditions that increase cardiomyocyte and nuclear strain, either by externally imposed stretch or by matrix-mediated tension. For patient-derived cardiac tissues, we contrast states of pathological hemodynamic load with mechanical unloading. We utilized human induced pluripotent stem cell-derived cardiomyocytes (hiPSC-CM), including patient PLN R14del, isogenic control and wild-type control cell lines, to investigate the response of these cells to cyclical mechanical stretch and differences in mechano-sensing when exposed to variations in substrate stiffness. Additionally, human living myocardial slices (LMS) of a PLN R14del patient are employed to study the effects of mechanical overload on nuclear alterations. Moreover, we analysed nuclear susceptibility to mechanical load in myocardial samples from patients with and without the PLN R14del variant by evaluating nuclear macro- and microstructure at the time of left ventricular assist device (LVAD) implantation, and at the time of heart transplantation. This integrative approach allows us to dissect the multi-scale impact of mechanical cues on nuclear structure and function in PLN R14del cardiomyopathy, from cellular mechanics to patient myocardium.

Our findings reveal that PLN R14del cardiomyocytes exhibited increased nuclear susceptibility to nuclear stress, as evidenced by abnormal calcium handling, altered nuclear morphology, and disrupted cytoskeletal integrity. These findings are consistently observed across cell models, patient LMS, and patient samples. Together, they establish a novel link between mechanical stress and nuclear dysfunction in PLN cardiomyopathy, identifying new insights into disease mechanism and highlighting potential therapeutic targets focused on nuclear mechanobiology.

## Methods

A detailed description of the methods is available in the supplemental material.

### Statistical analysis

Statistical analysis was performed using GraphPad Prism version 10.4.1, RStudio (RStudio Team), and R 3.5.0 (The R Foundation). One-way ANOVA was used for comparisons across multiple groups, with Tukey’s post-hoc test applied to identify significant differences between groups. For analyses involving repeated measures, a repeated-measures ANOVA was performed, followed by Tukey’s multiple comparisons test. A P-value of less than 0.05 was considered statistically significant. Values are presented as mean ± SD. For all immunofluorescence experiments, N represents experimental repeats, whereby each repeat is an average of 2-3 experimental replicates. Additionally, 5 random images were taken of each replicate, and the average was calculated as the value of that replicate. For Z-stack images, 3 random images were taken per replicate.

## Results

### PLN R14del results in nuclear abnormalities and calcium handling

The PLN R14del pathogenic variant has been primarily studied for its effects on SR calcium handling. However, altered SR calcium handling alone cannot fully explain the observed clinical heterogeneity in PLN R14del cardiomyopathy patients or asymptomatic carriers. Some limited evidence hints at the role of PLN in the nuclear envelope^33, 34^, but the effect of the R14del genetic variant on nuclear function remains underexplored. One previous report noted heavily wrinkled nuclei as a side observation in PLN R14del patients.^39^

Consistently, cardiomyocytes in the hearts of PLN R14del patients exhibit irregular nuclear shapes with invaginations, as shown in representative electron microscopy images of cardiac tissue (Figure 1A). These morphological changes are indicative of nuclear stress^40^ and suggest potential nuclear dysfunction in PLN R14del cardiomyopathy. To confirm the nuclear presence of PLN, we isolated nuclei from both cardiac tissue and hiPSC-CMs with PLN WT and PLN R14del. The purity of the nuclear fractions was validated by the absence of cytosolic and SR markers, GAPDH, and sarcalumenin, respectively. Western blot analysis demonstrated PLN expression in all nuclear fractions, except for the negative control, HeLa cells (Figure 1B). Immunofluorescence staining further confirmed PLN localisation in the nuclear envelope in PLN WT and PLN R14del tissue (Figure 1C), supporting the hypothesis that PLN is present and potentially plays a role in the nucleus. To assess the functional impact of the PLN R14del on nuclear calcium dynamics, we analysed cytosolic and nuclear calcium measurements of matured PLN WT and R14del hiPSC-CMs using DRAQ5™ segmentation (Figure S1), as indicated by the average calcium transient (Figure 1D). PLN R14del hiPSC-CMs exhibited an aberrant SR calcium handling, as indicated by the lower intrinsic beating rate (Figure 1E), prolonged beating rate-corrected calcium transient decay duration (CTD_90_; Figure 1F), and an increased calcium amplitude (Figure 1G). Additionally, nuclear CTD_90,_ as well as CTD_20_ and CTD_50_ (Figure S2A and S2B), were prolonged compared to the cytosolic counterpart in both PLN WT and R14del hiPSC-CMs, consistent with previous reports.^41^ Rise time was significantly prolonged in PLN R14del compared to PLN WT in both compartments, and slower in the nuclear compared to the cytosolic compartment in both genotypes (Supplementary Figure S2C). These findings collectively suggest that the previously reported disruption of SR function extends to nuclear calcium handling, further emphasizing the altered calcium dynamics in R14del hiPSC-CMs.

**Figure 1:**
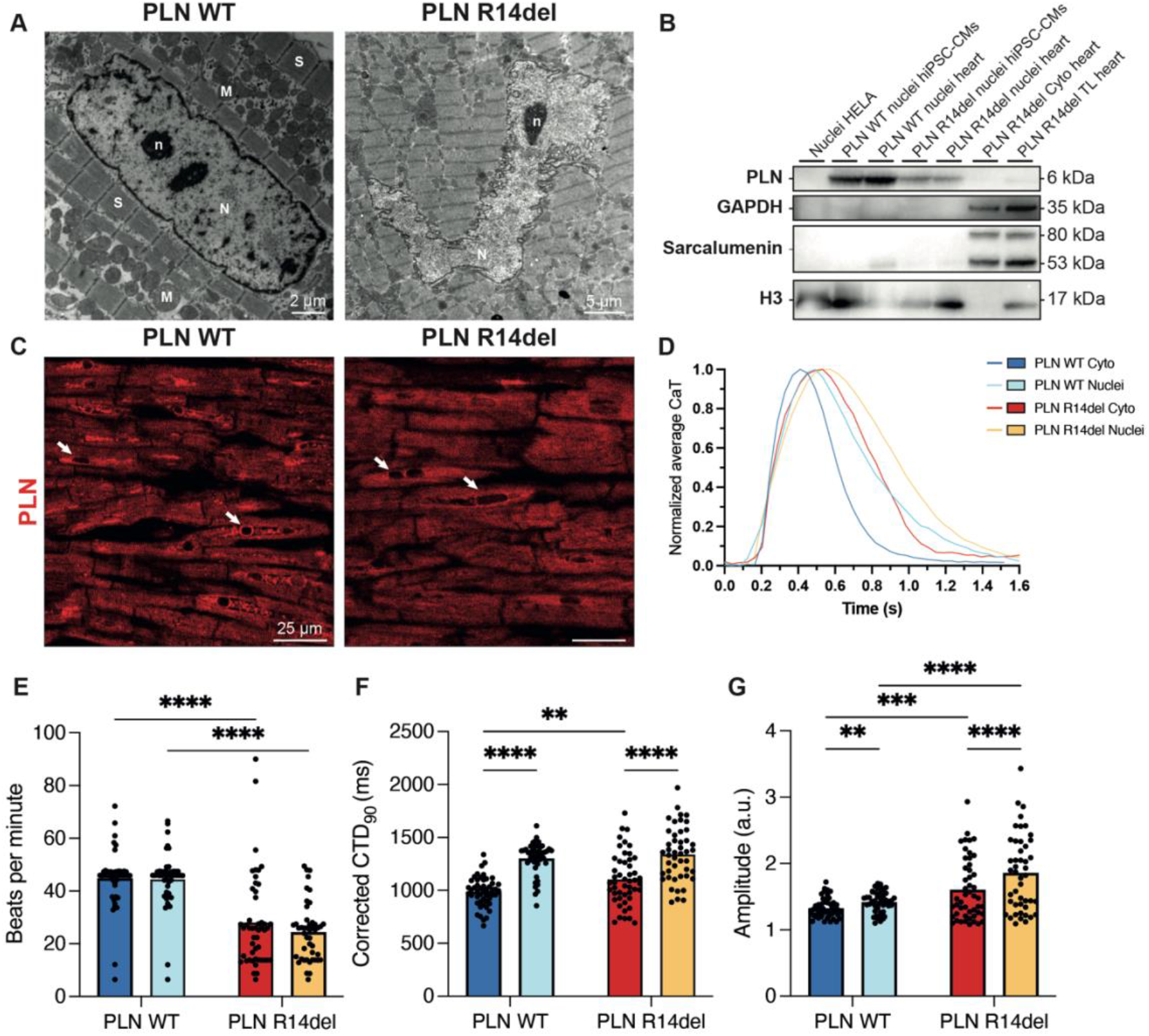
Stressed nuclei and impaired nuclear calcium handling in PLN R14del cardiomyocytes. A) Representative TEM images of cardiac tissue nuclei from a healthy control (PLN WT) and a PLN R14del patient. The nucleus (N) in healthy cardiac tissue displays an oval shape with a well-organized nucleolus (n), surrounded by sarcomeres (S) and mitochondria (M). In contrast, PLN R14del nuclei exhibit an irregular shape and invaginations, indicative of nuclear stress. hiPSC-CMs and cardiac tissue with PLN WT and PLN R14del show PLN protein expression in isolated nuclei (B) and in the nuclear envelope (C). Nuclear isolation is confirmed by the absence of sarcalumenin and GAPDH, markers for sarcoplasmic reticulum and cytosolic fractions, respectively, and the presence of H3 as a marker for nuclear fractions. D) Average calcium transients of nuclear and cytosolic compartments in matured PLN WT and R14del hiPSC-CMs reveal aberrant nuclear calcium dynamics. PLN R14del hiPSC-CMs show a lower beating rate (E), a prolongation of cytosolic calcium transient duration (F), and an elevated calcium amplitude, which is most pronounced in the nuclear compartment (G). (E-G: PLN WT n=50; PLN R14del n=50).

### PLN R14del hiPSC-CMs exhibit strain avoidance under cyclic mechanical stretch, but this cannot prevent cellular and nuclear damage

Previous work has demonstrated that nuclear calcium handling is an essential component of the strain avoidance response^37^, which remodels the cell to orient perpendicular to the direction of cyclic strain to protect the nucleus against stretch-induced damage. To study if this mechanism is affected in PLN R14del, we subjected PLN WT, R14del, and isogenic control (PLN corr) hiPSC-CMs to continuous cyclic uniaxial stretch (4%, 1 Hz) for 0, 30 min, 6 hrs, 24 hrs, and 48 hrs and evaluated nuclear injury, nuclear orientation, cytoskeletal dynamics, and cellular injury.

In PLN R14del hiPSC-CMs, we observed a significant increase in the nuclear injury, as assessed by the increased expression of the nuclear injury marker PML-NBs^42^, starting at 30 minutes and peaking at 6 hours of cyclic stretch (Figure 2A and 2B). This pattern was not observed in PLN WT and PLN corr hiPSC-CMs. Notably, after 24 hours of cyclic stretch, the nuclei of PLN R14del hiPSC-CMs reoriented to approximately 30° to the applied strain axis, coinciding with the reduction in the presence of PML-NBs (Figure 2C). Interestingly, in R14del hiPSC-CMs, F-actin fibres also remodeled approximately 30° away from the applied strain within 48 hours ^37^(Figure 2D, 2F, Figure S3A, and S3B). This effect was not observed in the PLN WT and PLN corr hiPSC-CMs. These findings suggest that PLN R14del hiPSC-CMs activate an adaptive strain avoidance response to protect the nucleus from mechanically induced damage.

**Figure 2:**
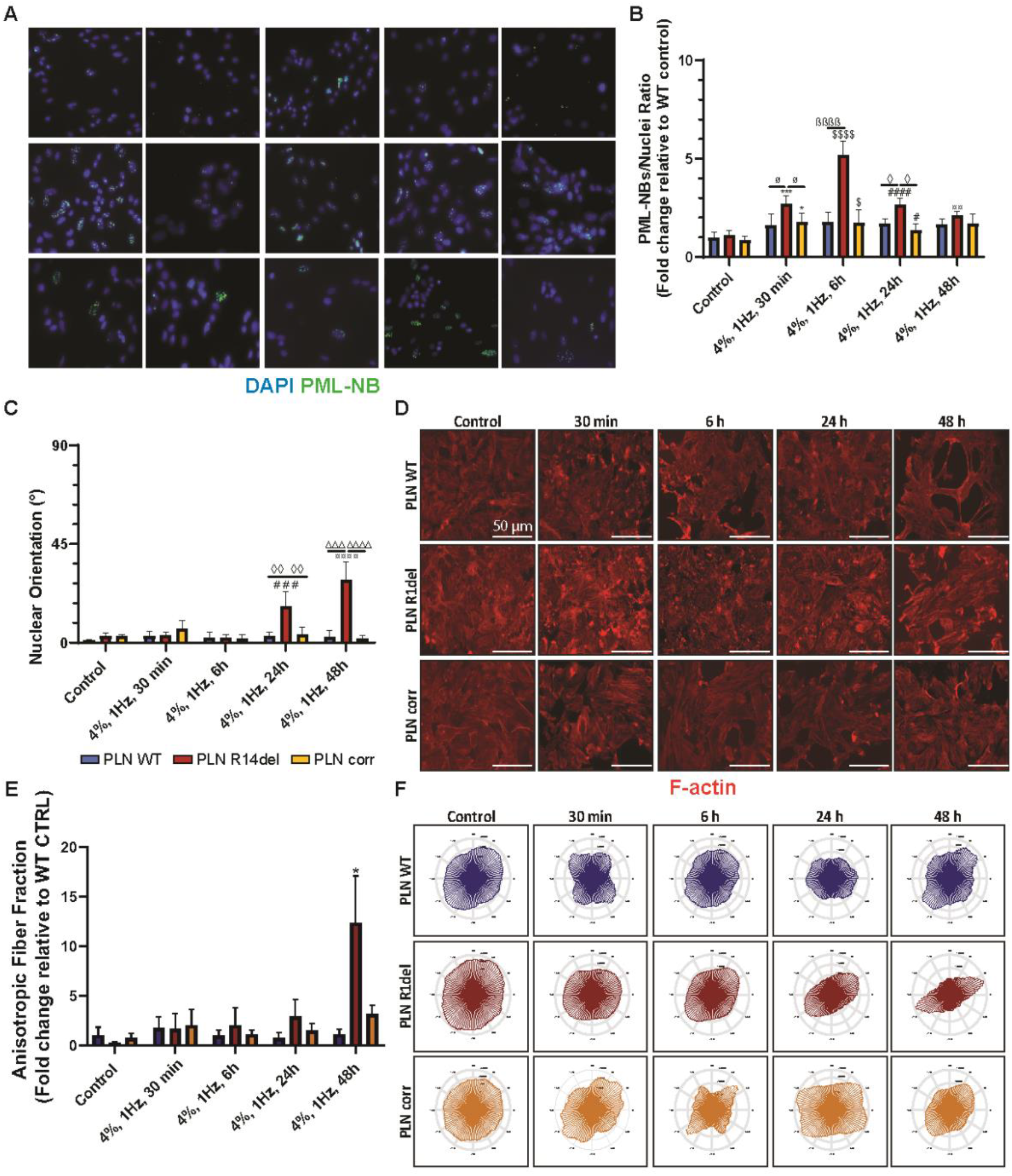
PLN R14del hiPSC-CMs are susceptible to mechanical load, leading to nuclear damage. A) Representative immunofluorescence images of DAPI-stained nuclei (blue) and PML-NBs (green) in PLN WT, PLN R14del, and PLN corr hiPSC-CMs in control conditions and after mechanical loading (4%, 1 Hz) applied in x-direction for 30 min, 6 h, 24 h, and 48 h. B) Quantification of the PML-NBs/nuclei ratio reveals a significant increase in PML-NBs in PLN R14del hiPSC-CMs under mechanical loading compared to PLN WT and PLN corr, starting at 30 min, peaking at 6 hours, and diminishing after 24 h. C) Quantification of nuclear orientation reveals a progressive reorientation toward 30° in PLN R14del hiPSC-CMs compared to PLN WT and PLN corr, observed after 24 hours of mechanical loading. Quantification of the anisotropic fibre fraction indicates alignment in PLN R14del hiPSC-CMs at 48 hours, compared to PLN WT and PLN corr D) Representative immunofluorescence images of F-actin (red) show fibre alignment in PLN R14del hiPSC-CMs under mechanical loading.(E), which is further corroborated by the polar plot (F). (B, D and E: PLN WT n=5; PLN R14del n=5; PLN corr n=5). Statistical significance is indicated by ø (30 min), β (6 h), ◊ (24 h), Δ (48 h), * (Control vs. 30 min), $ (Control vs. 6 h), # (Control vs. 24 h), and ¤ (Control vs. 48 h).

Next, we determined whether the strain avoidance response contributes to the ability of PLN R14del hiPSC-CMs to mitigate damage under stretch conditions. To assess this, we performed a temporal analysis of cellular apoptosis using TUNEL staining (Figure 3A). In PLN R14del hiPSC-CMs, TUNEL^+^ cells became increasingly prevalent at 24 hours and persisted until 48 hours after stretch initiation, whereas PLN WT and PLN corr hiPSC-CMs displayed minimal TUNEL positivity (Figure 3B). This heightened apoptosis was further corroborated by cleaved Caspase-3 (actCasp3) staining (Figure S4A and S4B). These findings suggest that although the PLN R14del hiPSC-CMs attempt to mitigate mechanically-induced injury by activating a strain avoidance response, this adaptation is insufficient to counteract the detrimental effects, culminating in increased cellular apoptosis. It must be noted that neither the PLN WT nor the PLN corr hiPSC-CMs demonstrated strain avoidance, although contrary to many adherent cell types cultured in 2D, this phenotype has been previously demonstrated^43^.

**Figure 3:**
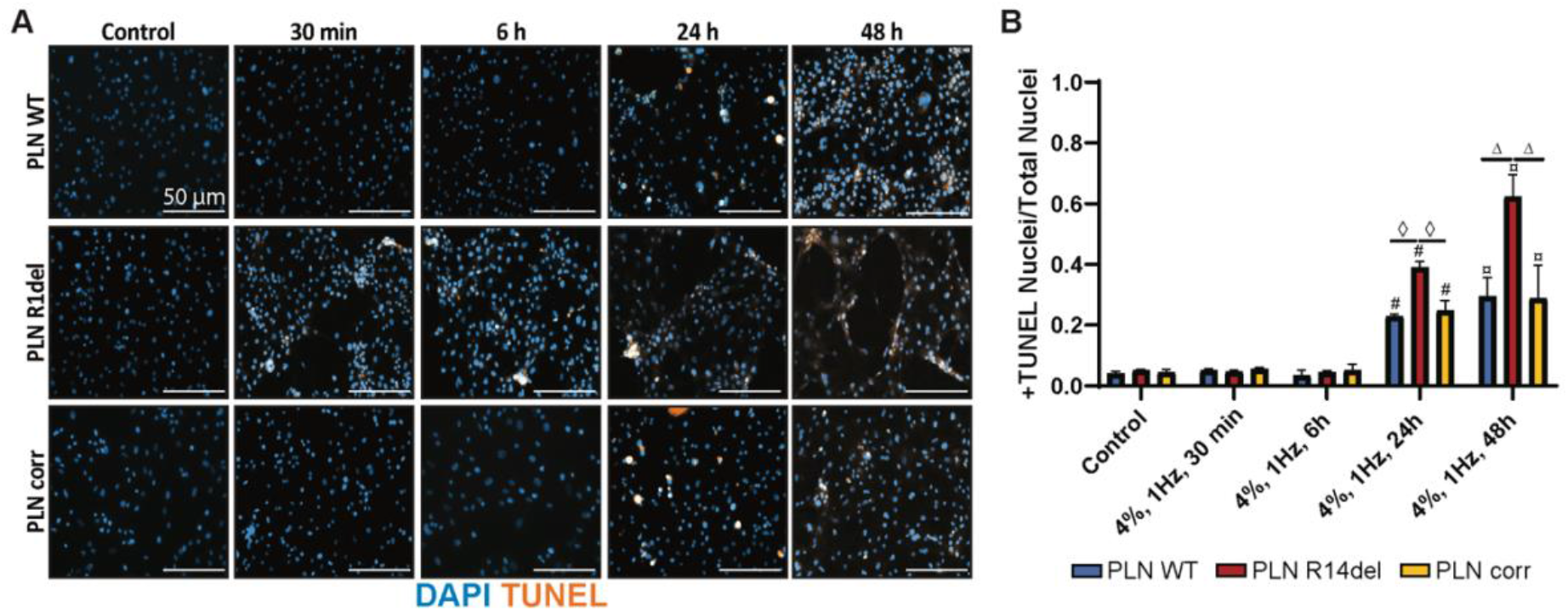
PLN R14del hiPSC-CMs undergo apoptosis in response to mechanical stress. A) Representative immunofluorescence images of TUNEL-positive nuclei (orange) and DAPI-stained nuclei (blue) in PLN WT, PLN R14del, and PLN corr hiPSC-CMs at control conditions and after mechanical loading (4%, 1 Hz) applied in x-direction for 30 min, 6 h, 24 h, and 48 h. B) Quantification of TUNEL-positive nuclei indicates elevated apoptosis in PLN R14del hiPSC-CMs, beginning at 24 hours and persisting after 48 hours of mechanical loading, compared to PLN WT and PLN corr (B: PLN WT n=5; PLN R14del n=5; PLN corr n=5). Statistical significance is indicated by ◊ (24 h), Δ (48 h), # (Control vs. 24 h), and ¤ (Control vs. 48 h).

### Nuclear and cytoskeletal abnormalities of hiPSC-CMs are evident at varying stiffness

In most adherent cell types, the actin cytoskeleton is reinforced in response to increased microenvironment stiffness^44, 45^, which compresses and protects the nucleus from mechanical stresses^46^. Although it is unknown whether this process also occurs in cardiomyocytes, it is known that cardiomyocytes produce a highly organised cytoskeletal structure when exposed to elevations in microenvironment stiffness (Figure S5)^47^. Since cardiomyocytes with the PLN R14del variant demonstrate an altered mechanotransduction when exposed to cyclic stretch, we further investigated if a similar response occurred to microenvironmental mechanical cues, specifically substrate stiffness. PLN WT and PLN R14del hiPSC-CMs were cultured on PAA hydrogels with physiological stiffness (10-20 kPa) comparable to the healthy myocardium^42, 48^ and pathological stiffness (65-85 kPa) comparable to a diseased heart.^49^ After 3 days of culture, cells were fixed and stained with DAPI and F-actin (Phalloidin 647N) to assess nuclear and cytoskeletal organisation (Figure 4A). Our analysis demonstrated that both PLN WT and PLN R14del hiPSC-CMs respond to increased microenvironment stiffness by enhancing actin fibres abundance in the cell (Figure 4B). By using z-stack imaging to evaluate the nuclear and actin morphology in 3D, no difference in the nuclear area was observed in PLN WT and PLN R14del hiPSC-CMs (Figure 4C and 4D) on different substrates. However, nuclear height was consistently greater in PLN R14del hiPSC-CMs compared to PLN WT, regardless of substrate stiffness (Figure 4E). Importantly, an inverse trend was observed for the height of cytoskeletal F-actin staining, which was lower in PLN R14del hiPSC-CMs under both stiffness conditions (Figure 4E). This suggests that nuclei in PLN R14del hiPSC-CMs are inadequately shielded by F-actin nuclear encapsulation, leaving them vulnerable to mechanical stresses. This vulnerability was supported by the increased expression of PML-NBs in the PLN R14del hiPSC-CMs (Figure 4F and 4G).

**Figure 4:**
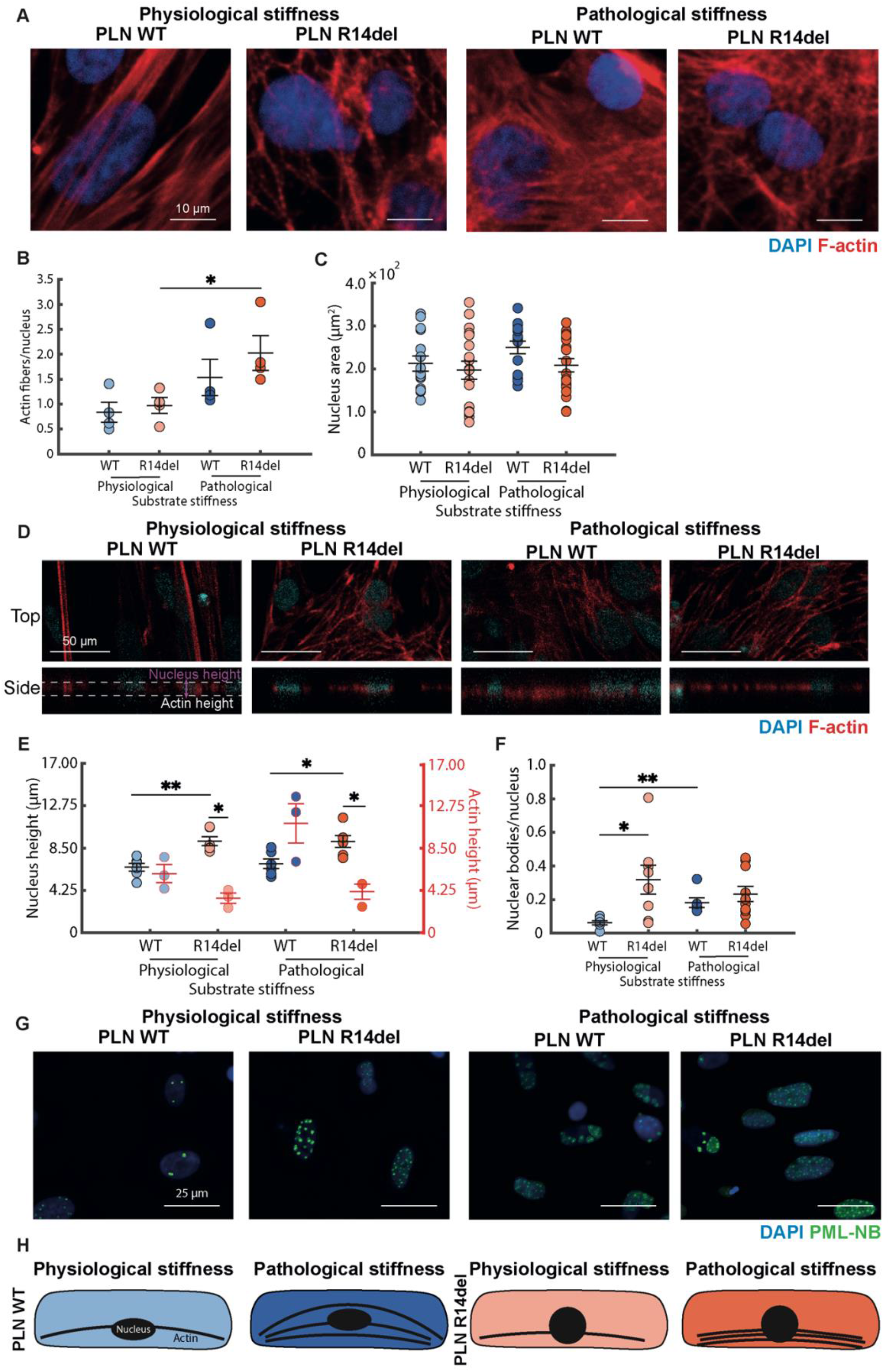
PLN R14del variant leads to abnormal nuclear and actin cytoskeletal structure in cardiomyocytes under the influence of microenvironment stiffness. A) Representative immunofluorescence images show DAPI (blue) and F-actin (red) organisation in PLN WT and PLN R14del hiPSC-CMs on substrates with physiological and pathological stiffness. B) Quantification of actin fibres per nucleus reveals a significant increase in PLN R14del hiPSC-CMs under pathological stiffness compared to PLN WT (PLN WT n=4; PLN R14del n=4). C) Quantification of the nuclear area shows no significant difference between PLN WT and PLN R14del under either stiffness condition. D) Representative z-stack immunofluorescence images of DAPI-stained nuclei (blue) and F-actin (red) in PLN WT and PLN R14del hiPSC-CMs cultured on substrates with physiological and pathological stiffness. The top and bottom row panels display the top and the side of the monolayers, respectively. E) nuclear height is significantly reduced in PLN R14del hiPSC-CMs under both physiological and pathological stiffness compared to PLN WT (C: PLN WT n=15; PLN R14del n=18, D: PLN WT n=6; PLN R14del n=6). F) Quantification of the PML-NBs/nuclei ratio shows a significant increase in PML-NBs in PLN R14del hiPSC-CMs compared to PLN WT under physiological stiffness, while this increase is only apparent for PLN WT under pathologic stiffness (10-20 kPa PLN WT n=6; 65-85 kPa PLN WT n=6; 10-20 kPa PLN R14del n=8, 65-85 kPa PLN R14del n=10). G) Representative immunofluorescence images of DAPI-stained nuclei (blue) and PML-NBs (green) in PLN WT, PLN R14del on substrates with physiological and pathological stiffness. H) Schematic representation of actin and nuclear structure of PLN WT and R14del hiPSC-CMs under physiological and pathological stiffness.

Collectively, these findings indicate that although all cells reinforce their actin cytoskeleton in response to increased substrate stiffness (Figure 4H), increased nuclear height coupled to reduced F-actin height (indicating the lack of actin cap formation) in PLN R14del hiPSC-CMs suggests ineffective nuclear encapsulation. This structural vulnerability may render their nuclei more susceptible to mechanical strain, consistent with the observed increase in nuclear PML-NB expression.

### PLN R14del variant impairs LINC-mediated mechanotransduction and YAP signalling

The LINC complex mechanically connects the actin cytoskeleton to the nucleus.^50^ We performed immunofluorescence staining for two key components of the LINC complex, Desmin and Nesprin-1, both implicated in the onset of cardiomyopathies.^51, 52^ While the overall intensity of Desmin in the cytosol was similar between PLN WT and R14del hiPSC-CMs (Figure 5A), its cellular distribution differed. On substrates with pathological stiffness, Desmin appeared diffusely in the cytoplasm of PLN WT hiPSC-CMs, whereas large perinuclear aggregates were observed in PLN R14del hiPSC-CMs (red arrows, Figure 5A to 5C). Similarly, the total intensity of Nesprin-1 was comparable in PLN WT and R14del hiPSC-CMs on physiological stiffness (Figure 5D). However, on pathological stiffness, Nesprin-1 intensity was markedly reduced in PLN R14del hiPSC-CMs, especially in the peri-nuclear region (Figure 5E and 5F), and thereby likely contributed to the aberrant actin structure observed.

**Figure 5:**
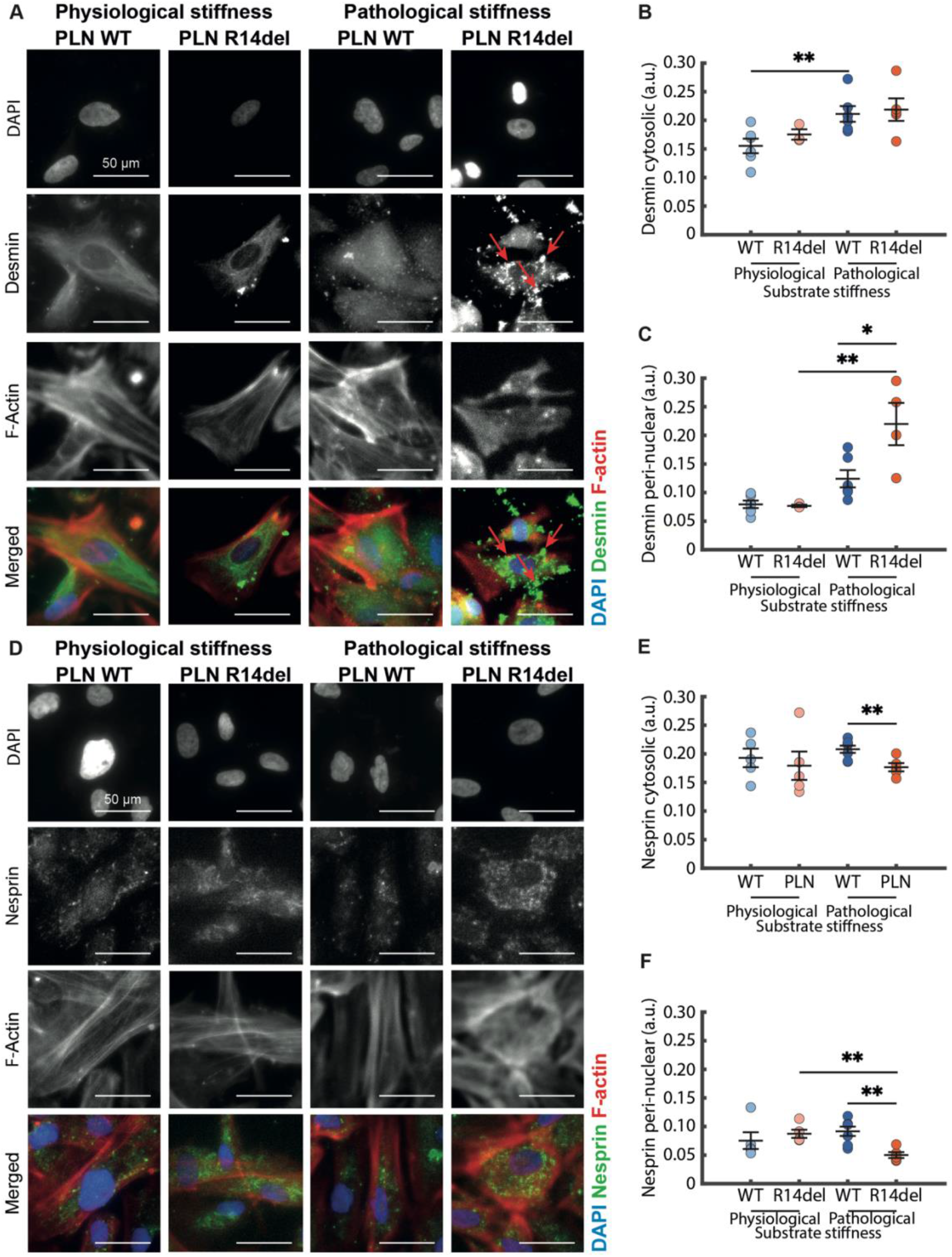
PLN R14del variant results in compromised mechanotransduction when exposed to pathological stiffness. A) Representative immunofluorescence images of DAPI-stained nuclei (blue), Desmin (green), and F-actin (red) in PLN WT and PLN R14del hiPSC-CMs cultured on substrates with physiological and pathological stiffness. Red arrows highlight Desmin abnormalities in PLN R14del hiPSC-CMs under pathological stiffness. Quantification of Desmin levels shows elevation of both cytosolic (B) and perinuclear Desmin (C) under pathological stiffness in PLN WT and R14del hiPSC-CMs, with perinuclear Desmin being more apparent in PLN R14del hiPSC-CMs (15 kPa PLN WT n=6; 65-85 kPa PLN WT n=6; 15 kPa PLN R14del n=3, 65-85 kPa PLN R14del n=5). D) Representative immunofluorescence images of DAPI-stained nuclei (blue), Nesprin-1 (green), and F-actin (red) under similar conditions. Quantification showed a lower cytosolic Nesprin-1 expression in PLN R14del hiPSC-CMs compared to WT under pathological stiffness (E), in line with peri-nuclear Nesprin-1 expression (F), which was already decreased from physiological to pathological stiffness for PLN R14del hiPSC-CMs (15 kPa PLN WT n=5; 65-85 kPa PLN WT n=7; 15 kPa PLN R14del n=5, 65-85 kPa PLN R14del n=5).

Since cardiomyocytes continuously generate mechanical stresses through both sarcomeric contraction and actomyosin-mediated cytoskeletal tension, we subsequently assessed whether the observed alterations in the LINC complex were associated with changes in cellular mechanical stress and force generation using traction force microscopy (Figure S6). Increasing substrate stiffness resulted in elevated diastolic and systolic mechanical stresses in both genotypes (Figure S6A-C). However, this response was substantially more pronounced in PLN R14del hiPSC-CMs, indicating increased intrinsic cellular tension under pathological stiffness conditions. We subsequently investigated whether this increase in mechanical stress translated into enhanced contractile performance. Consistent with adaptive mechanobiological responses previously reported in healthy cardiomyocytes, PLN WT hiPSC-CMs responded to pathological stiffness with an increase in effective contraction (Figure S6D and S6E). In contrast, PLN R14del hiPSC-CMs failed to show a corresponding increase in effective contraction despite generating substantial mechanical stresses.

Thus, whereas PLN WT cardiomyocytes exhibited increased effective contraction in response to pathological stiffness, PLN R14del cardiomyocytes generated elevated mechanical stresses without a corresponding improvement in contractile performance. These findings suggest a decoupling of mechanical stress generation from productive contractile function in PLN R14del cardiomyocytes, consistent with impaired mechanotransduction and force transmission.

Since an intact LINC complex is critical for mechanotransduction via YAP signalling^44, 45, 53, 54^, we further assessed the nuclear localisation of the active, non-phosphorylated YAP on pathological stiffness (Figure S7A). As expected, PLN WT hiPSC-CMs demonstrated mechanobiological adaptation to pathological stiffness conditions by increased nuclear localisation of YAP. In contrast, PLN R14del hiPSC-CMs failed to increase nuclear YAP localisation under the same conditions (Figure S7B).

These findings suggest that the PLN R14del variant disrupts the structural integrity of the LINC complex and impairs YAP-mediated mechanotransduction, exacerbating nuclear and cytoskeletal dysregulation under pathological stiffness.

### PLN R14del cardiomyopathy patient tissues corroborate nuclear abnormalities under mechanical stress

To assess the clinical relevance and confirm the observed nuclear and cytoskeletal abnormalities in hiPSC-CMs, we analysed cardiac tissue from PLN R14del and DCM patients of suspected but unidentified genetic origin, and healthy controls. A summary of baseline characteristics of patients is provided in Table 2. For PLN R14del and DCM patient tissues, nuclear morphology was assessed at two distinct time points for each patient: during mechanical loaded state at the time of LVAD implantation and the mechanically unloaded state at heart transplantation. Immunofluorescence staining of PCM-1 and DAPI revealed nuclear abnormalities in loaded PLN R14del patients relative to their unloaded counterparts and DCM patients (Figure 6A). The nuclear area was comparable between healthy controls and other DCM patients across both time points, with trends varying per individual. PLN R14del patients in a loaded state consistently exhibited an increased nuclear area, which normalized upon unloading (Figure 6B). Similarly, nuclear circularity was reduced in loaded PLN R14del patients compared to healthy controls and DCM patients, while unloading restored circularity (Figure 6C). Conversely, nuclear eccentricity was higher under mechanical load and decreased upon unloading in PLN R14del patients, which was not seen in the DCM group (Figure 6D). To further investigate nuclear ultrastructure, transmission electron microscopy (TEM) was used to assess the regularity of nuclear integrity by measuring the ratio of the nuclear perimeter with and without invaginations (Figure 6E). Invaginations, indicative of nuclear stress, were more pronounced in loaded PLN R14del patients, which decreased upon unloading (Figure 6F).

**Figure 6:**
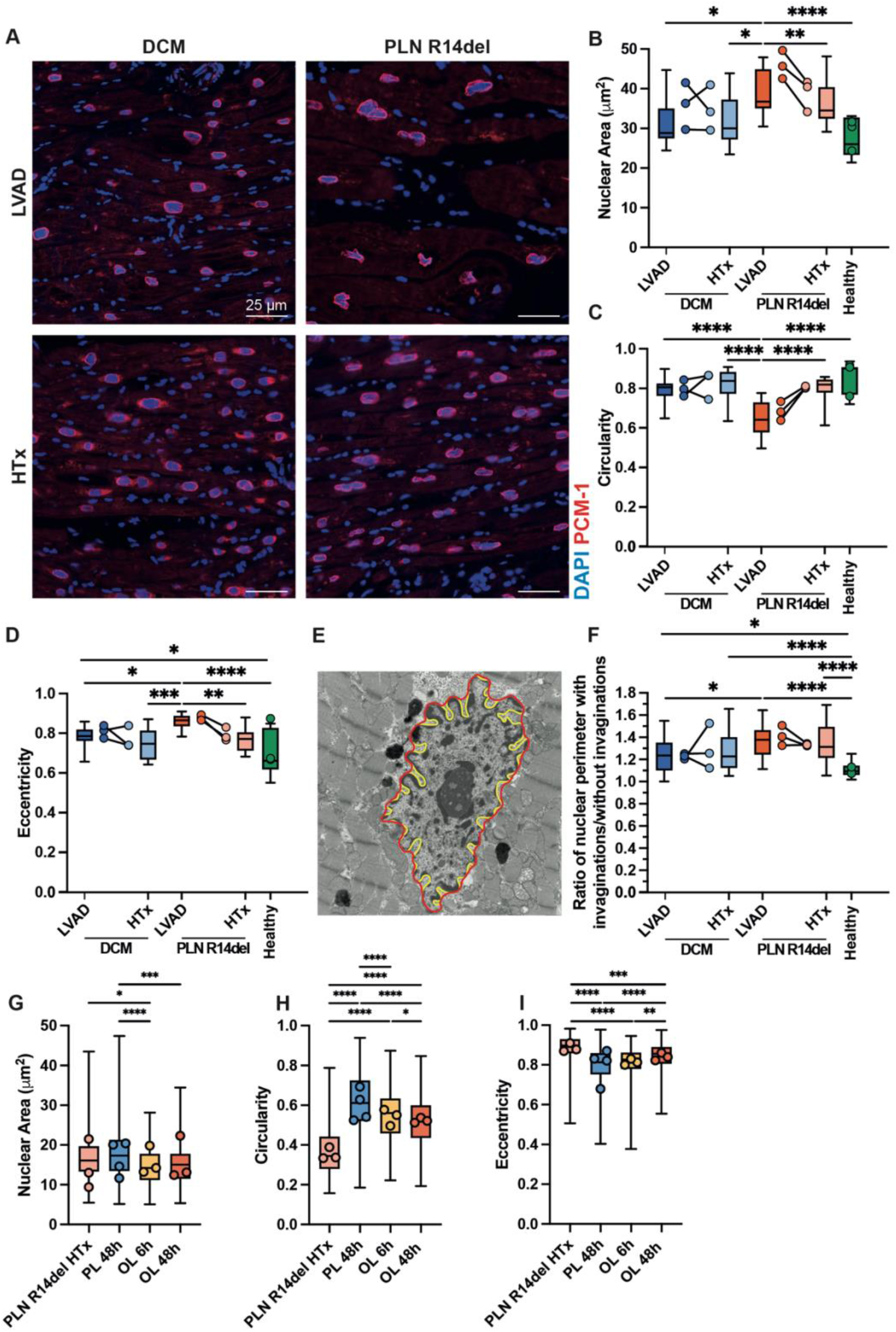
PLN R14del patients exhibit nuclear abnormalities under pathological loading conditions. A) Representative immunofluorescence images of DAPI-stained nuclei (blue) and PCM-1 (red) in cardiac tissue from PLN R14del and DCM patients under mechanical load at the time of LVAD implantation and unloaded condition at the time of HTx. B) Quantification of the nuclear area showed that in a diseased state, the nuclear area increases, which is exacerbated in the loaded state of patients harboring the PLN R14del variant. Similarly, circularity (C) and eccentricity (D) reveal that PLN R14del nuclei have a decreased circularity and increased eccentricity compared to the loaded DCM patients (B-D: DCM N=3, LVAD, n=1874; HTx, n=1661; PLN R14del, N=3, LVAD, n=1176; HTx, n=1708; Healthy, N=3, n=1808). E) TEM image showing invaginations in the nucleus, with the perimeter with and without invaginations. F) The ratio of invagination is increased in loaded PLN R14del condition and normalizes upon unloading (DCM N=3, LVAD n=28; HTx n=27; PLN R14del N=3, LVAD n=21; HTx n=27; Healthy N=3, n=42). Analysis of nuclear morphology in LMS derived from a PLN R14del patient subjected to overloading (OL) for 6 hours or 48 hours and physiological loading (PL) for 48 hours, compared to the fresh sample at HTx. Quantification of nuclear area (G), circularity (H), and eccentricity (I) demonstrates that normalization of nuclear morphology is observed under physiological loading conditions, which is inhibited by the overloading (HTx N=3, n=131; PL 48h N=4, n= 543; OL 6h N=3, n=499; OL 48h N=3, n= 353).

**Table 1.** Concentrations of acrylamide and bis-acrylamide (% w/v) to prepare PAA substrates of 10-20 and 65-85 kPa.

| Stiffness | 10-20 kPa | 65-85 kPa |
| --- | --- | --- |
| Acrylamide (% w/v) | 7.55 | 12 |
| Bis-acrylamide (% w/v) | 0.12 | 0.37 |

**Table 2:**
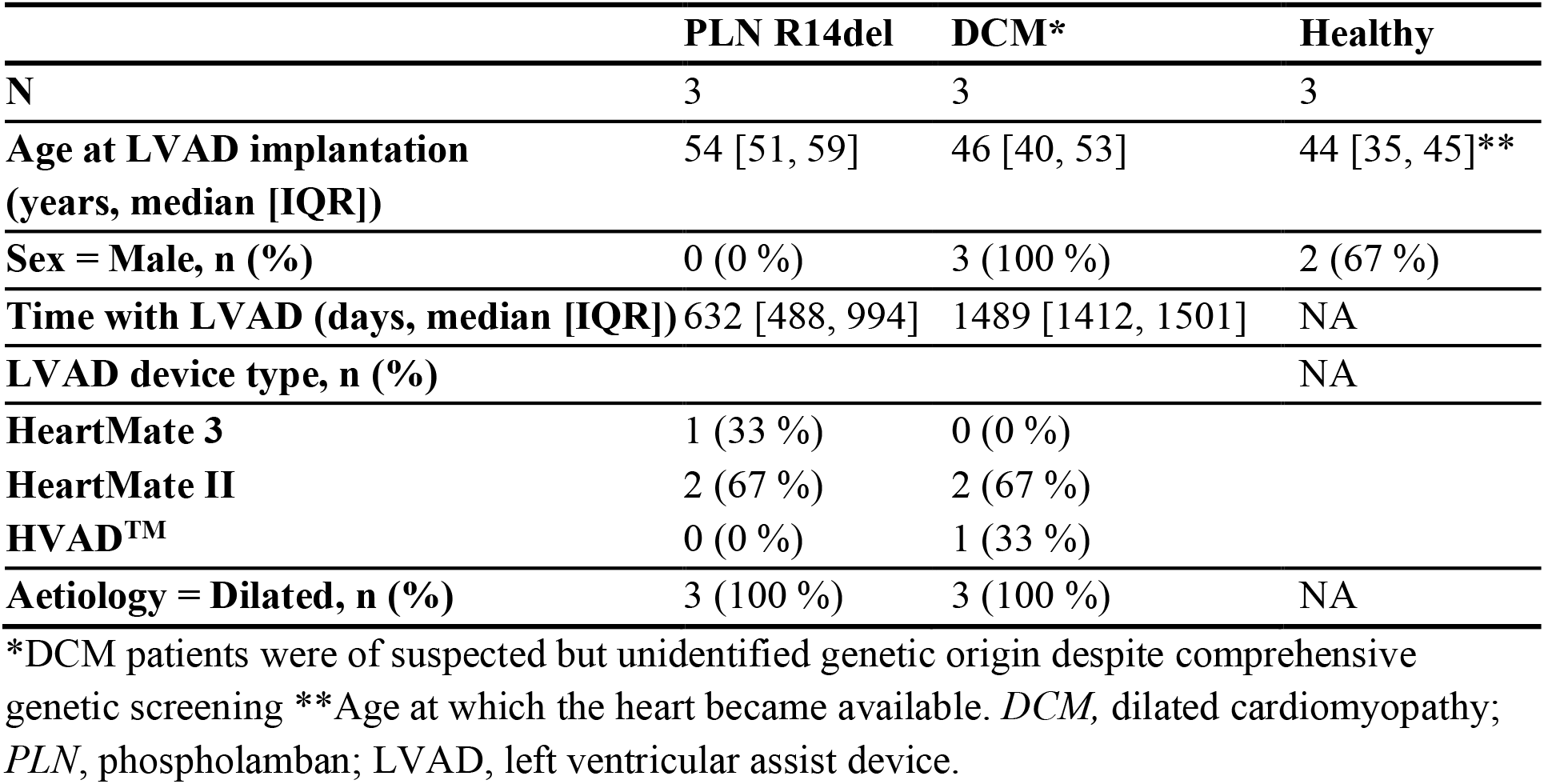
Baseline characteristics of patients’ biopsies used for nuclear analysis at LVAD.

To directly assess acute mechanosensing defects in human PLN R14del cardiac tissue, within a timeframe comparable to hiPSC-CMs experiments, LMS (N=11) were generated from the left ventricular free wall of a 52-year-old male PLN R14del patient at the time of heart explantation. LMS were subjected to either quasistatic overloading (OL), by stretching to 29% of the original slice length or maintained under physiological loading (PL) conditions^55^. Nuclear morphologic analysis revealed that physiological loading normalized nuclear morphology in PLN R14del LMS, reflected by a shift towards increased nuclear area (Figure 6G), increased circularity (Figure 6H), and reduced eccentricity (Figure 6I) compared to freshly explanted tissue. In contrast, mechanical overloading prevented this normalization, recapitulating the nuclear abnormalities observed in loaded PLN R14del patient hearts at the time of LVAD implantation.

These findings demonstrate that the nuclear abnormalities observed in PLN R14del hiPSC-CMs *in vitro* are recapitulated in patient cardiac tissue, underscoring a specific vulnerability of PLN R14del cardiomyocytes to mechanical loading. This highlights nuclear mechanotransduction dysfunction as a distinctive feature of PLN R14del cardiomyopathy pathogenesis, not observed to the same extent in DCM.

## Discussion

While the role of PLN R14del in SR calcium handling has been extensively studied, its implications for mechanotransduction and nuclear calcium signalling in cardiomyocytes remain largely unexplored. In this study, we demonstrate that the disruption of nuclear calcium handling by the PLN R14del variant renders cardiomyocytes vulnerable to mechanical stress, manifesting as nuclear and cytoskeletal abnormalities that represent a previously unrecognized pathological feature of PLN R14del cardiomyopathy.

Our findings reveal impaired nuclear calcium handling in PLN R14del hiPSC-CMs, in line with the known dependency of nuclear calcium on SERCA-mediated cytosolic calcium dynamics.^56, 57^ Since nuclear calcium is a key regulator of nuclear remodelling under mechanical stress^37^, our results highlight how PLN R14del cardiomyocytes exhibit heightened vulnerability to mechanical stimuli, leading to nuclear damage that, if persistent, culminates in cellular apoptosis.

While PLN R14del in the SR is predominantly investigated by its disruptive role on SERCA^58–60^, nuclear calcium transients mediated by inositol 1,4,5-trisphosphate receptors (IP3Rs) have been shown to activate transcriptional regulators involved in cardiac remodelling, including Nuclear Factor of Activated T-cells (NFAT), Ca²⁺/Calmodulin-dependent Protein Kinase II (CaMKII), and Myocyte Enhancer Factor 2 (MEF2)-dependent pathways.^61^ Given the impaired nuclear calcium dynamics observed in PLN R14del cardiomyocytes, altered IP3R-mediated calcium release may further contribute to defective nuclear signalling and mechanoadaptation. Moreover, mechanical stimuli themselves can modulate phospholipase C-IP3 signalling, raising the possibility that abnormal mechanotransduction and nuclear calcium dysregulation form a reinforcing pathological feedback loop in PLN R14del cardiomyopathy.^62^ While these various nuclear calcium mechanisms were not directly assessed in the present study, future investigations examining the spatial organization and potential disrupted function could provide additional mechanistic insight into how altered calcium signalling contributes to nuclear dysfunction and disease progression.

An important question arising from our findings is whether the observed defects in nuclear calcium handling are a cause or a consequence of altered nuclear morphology. Although our data do not allow us to establish the directionality of this relationship, accumulating evidence suggests that nuclear calcium signalling and nuclear architecture are closely interconnected. On the one hand, altered nuclear calcium dynamics may contribute directly to nuclear fragility. Nuclear calcium has been implicated in chromatin remodelling and adaptive nuclear responses to mechanical stress, processes that can protect the nucleus from excessive mechanical strain and damage.^33, 34, 37^ Impaired nuclear calcium signalling could therefore compromise these adaptive responses, rendering PLN R14del nuclei more susceptible to deformation and damage. On the other hand, structural abnormalities of the nuclear envelope may themselves disrupt calcium homeostasis by altering the organization of nuclear envelope-associated calcium-handling proteins and the calcium exchange between the nucleoplasm and the surrounding sarcoplasmic reticulum. Related nuclear envelope abnormalities and cellular dysfunction have been described in laminopathies and arrhythmogenic cardiomyopathies^63–65^, supporting the possibility that these processes may reinforce one another.^66^

Our findings are consistent with this concept, in which altered nuclear calcium signalling and nuclear structural abnormalities form a self-amplifying cycle that may ultimately converge on impaired mechanotransduction and cellular dysfunction in PLN R14del cardiomyopathy.

In line with this, in response to cyclic mechanical stretch, PLN R14del hiPSC-CMs attempt to compensate via a strain-avoidance mechanism^37^, including nuclear and F-actin reorientation as an adaptive response to redistribute mechanical load. This appears to be the first report of such strain-avoidant behaviour in immature cardiomyocytes, suggesting an innate capacity of PLN R14del cardiomyocytes to counteract their mechanical vulnerability even at early developmental stages.

Moreover, exposure to pathological substrate stiffness induces pronounced nuclear and cytoskeletal defects in PLN R14del hiPSC-CMs. Mechanistically, this is associated with elevated intrinsic cellular tension, impaired contractile function, and mechanosensory failure, as evidenced by the accumulation of LINC complex components (Desmin and Nesprin-1) and the inability of YAP, a key mechanotransduction effector, to translocate to the nucleus. YAP signalling is known to regulate cardiac mechanical integrity, contractility, calcium handling, and neonatal regeneration.^67–69^ In WT cardiomyocytes, increased matrix stiffness triggers YAP nuclear translocation via mechanosensitive nuclear pores, a process dependent on the transmission of actomyosin-generated tension through the LINC complex.^44^ In contrast, PLN R14del cardiomyocytes fail to activate YAP under the same mechanical conditions, supporting the idea that their mechanotransduction failure originates from compromised cytoskeletal and nuclear coupling.

The observation that PLN R14del cardiomyocytes generate high mechanical stresses despite reduced effective contraction suggests an inefficient mechanical state in which increased mechanical stress is not effectively translated into productive contractile output. The prolonged Ca²⁺ transient decay observed in PLN R14del hiPSC-CMs suggests that impaired Ca²⁺ removal contributes to this mechanical inefficiency by impairing myofilament relaxation between contractile cycles. Notably, the elevated Ca²⁺ amplitude would be expected to enhance rather than reduce effective force generation, further supporting the interpretation that the reduced effective contraction reflects impaired force transmission rather than a primary deficit in Ca²⁺- driven myofilament activation. Additionally, the observed disruption of the LINC complex and impaired nuclear YAP translocation, suggest alterations in the mechanosensing machinery involved in coordinating cellular adaptation to increased mechanical load. Consequently, PLN R14del cardiomyocytes appear to exhibit elevated mechanical stress coupled with impaired mechanosensory signalling, a combination that may contribute to progressive nuclear damage and cellular dysfunction.The integrity of the LINC complex appears particularly sensitive to cytosolic calcium levels, as its assembly is promoted by low calcium ion concentrations.^70, 71^ PLN R14del has been associated with high cytosolic calcium levels^6^, which could impair LINC complex formation, further weakening force transmission to the nucleus. Although the precise impact of the PLN R14del variant on calcium cycling is debated, the convergence of calcium dysregulation, cytoskeletal disorganisation, and YAP inactivation strongly implicates a mechanotransduction defect as a driver of cellular dysfunction in PLN R14del cardiomyopathy.

Importantly, these *in vitro* findings could be further validated by patient-derived materials. In cardiac tissues from PLN R14del patients, nuclear abnormalities, such as invaginations and irregular shapes, were prominently observed under mechanical loading conditions (*i.e.*, at the time of LVAD) and reversed upon mechanical unloading at the time of transplantation. This suggests that nuclear structure in PLN R14del cardiomyocytes is dynamically responsive to mechanical cues and that mechanical stress acts as a modifiable disease driver. Consistent with this, LMS generated from PLN R14del patients demonstrated that physiological loading was sufficient to normalize nuclear morphology, while overloading prevented this nuclear adaptation, mirroring the loaded state observed in patients at LVAD implantation.

Together, these findings emphasize a central role for nuclear mechanotransduction dysfunction in the pathogenesis of PLN R14del cardiomyopathy. They also establish a causal link between external mechanical stimuli and nuclear failure, demonstrating that aberrant mechanotransduction is not just a consequence but a plausible driver of disease in PLN R14del hearts.

### Aberrant mechanotransduction in cardiomyopathies

Nuclear defects are increasingly recognized as substates for arrhythmogenic cardiomyopathy (ACM), especially in variants affecting *PKP2*^64^, *LMNA*^65^, and *LEMD2*^63^. These genes are crucial for maintaining nuclear integrity and cell-cell connections, which directly link to mechanotransduction. Although PLN has been traditionally viewed as a cytosolic calcium regulator, our findings position PLN R14del within this framework, suggesting that, beyond calcium handling, PLN plays a previously underappreciated role in mechanotransduction, likely through its interactions at the nuclear envelope and its indirect influence on the LINC complex.

The PLN R14del variant appears to amplify mechanotransduction failure, combining aberrant calcium signalling with compromised nuclear-cytoskeletal coupling to drive cardiomyopathy progression. This mechanobiological impairment may be especially relevant for explaining the heterogeneous clinical manifestation of PLN R14del and highlights mechanosignalling as a converging point of diverse cellular dysfunctions. While these effects are pronounced in PLN R14del, altered intracellular calcium dynamics are a hallmark of cardiac remodelling and failure more broadly, suggesting that mechanotransduction failure may be a shared feature of cardiomyopathies and an attractive therapeutic target

### Mechanical stress in the progression of PLN R14del cardiomyopathy

Mechanical stress, resulting from intense physical activity (especially sports-related mechanical stress), is known to influence disease progression in ACM.^72, 73^ However, in PLN R14del patients, moderate physical activity does not appear to worsen disease progression^74^, although the effects of extreme exertion remain unclear and the study was likely underpowered due to a large number of unaffected relatives. Similarly, chronic stress, which also elevates cardiac mechanical load, has not been definitively linked to disease progression.^75^ It must be noted that these clinical studies rely on patient-reported questionnaires, which are inherently prone to biases and limitations. In contrast, preclinical models support a role for mechanical stress in PLN R14del pathology. Mice harbouring the human PLN R14del variant exhibit worsened arrhythmogenic phenotypes with exercise, reinforcing a causative link between mechanical loading and disease exacerbation.^76^

Anatomically, PLN R14del cardiomyopathy often originates in the apex of the heart^77^, a region that experiences non-uniform mechanical stress due to its unique mechanical myofibre orientation and deformation dynamics.^78–81^ Pre-symptomatic carriers with early apical deformation patterns are at significantly increased risk of arrhythmias within 3 years.^77^ This regional vulnerability underscores how localised mechanical stress may act as a trigger or amplifier of PLN R14del pathology, further implicating mechanotransduction failure as an important determinant of disease progression.

Taken together, these insights suggest that mechanical stress, tissue stiffness, and mechanoresponsiveness may collectively influence disease severity and the clinical presentation. They also highlight the need for careful evaluation of exercise guidelines for PLN R14del patients, balancing the potential risks of exacerbating cardiac dysfunction with the known benefits of physical activity.

## Study limitations

Exploring whether similar nuclear abnormalities occur in other PLN variants, such as the cardiomyopathy-causing PLN knock-out variant (PLN c.117T>A, p.Leu39*)^82^, would be essential to determine if nuclear dysfunction is a universal feature of all PLN-related cardiomyopathies or unique to the R14del. While our study integrates patient-derived hiPSC-CMs, including an isogenic control, *ex vivo* LMS, and patient-derived tissues, the tested patient cohort used for tissue analysis remains small. Future studies in larger cohorts with diverse clinical manifestations, accounting for sex-specific differences and the influence of LVAD support duration, will be critical to establish a mechanobiological risk profile for PLN R14del cardiomyopathy. Additionally, the observed reversibility of the nuclear abnormalities after LVAD-induced unloading suggests therapeutic potential, and a mechanobiological risk profile could identify which patients might benefit most from LVAD-mediated unloading. Together, these studies could advance our understanding of nuclear mechanotransduction in cardiac health and disease, paving the way for novel therapeutic strategies.

## Conclusion

This study signals a role for impaired nuclear mechanotransduction in the pathogenesis of PLN R14del cardiomyopathy, linking mechanical stress, cytoskeletal disorganisation, LINC complex dysfunction, and calcium signalling abnormalities to nuclear damage and cell death. Our findings highlight how YAP inactivation, resulting from disrupted cytoskeletal tension transmission, further compromises the cellular mechanotransduction under pathological stiffness. By uncovering this convergence between calcium and mechanical signalling pathways, we provide a more integrated view of PLN R14del pathogenesis. Importantly, this mechanistic insight sheds new light on the clinical heterogeneity observed among PLN R14del carriers. This work paves the way for mechanobiologically based interventions aimed at restoring nuclear integrity and mechanoresponsiveness, offering a new direction for precision therapy in genetic cardiomyopathies.

## Acknowledgments

The authors thank Christian Snijders Blok, Inge Dokter, Mark Daniels, and Petra van der Kraak for their valuable technical expertise. We gratefully acknowledge Tomo Saric (Uniklinik Köln) for providing the human iPSC control line and Prof. Mark Mercola (Stanford University) for sharing the D4iR14del and C31iCTR lines. We also acknowledge the Cell Microscopy Core (CMC) of the Center for Molecular Medicine, UMC Utrecht, for their microscopy training and support. Finally, we would like to thank the close collaboration between cardiothoracic surgeons (CTC), cardiologists, heart failure nurses, the OR team, and VAD coordinators of the UMC Utrecht for their valuable assistance in procuring fresh cardiac tissue.

## Data Availability

The authors confirm that the data supporting the findings of this study are available within the article and its supplementary materials. The raw data that support the findings of this study can be made available on request from the corresponding authors CVCB, JPGS, and AvdP.

## Author Contributions

JSAvdG, AvdP, VC, JPGS, and CVCB, conceptualized and designed the study. JSAvdG, AvdP, LH, AA, JvA, and MAMEvR participated in conducting the experiments and data acquisition. JSAvdG, AvdP, LH, AA, JvA, MAMEvR, WBvH, ASJMtR, AvM, VSP, PAD, ZL, VC, PGS, and CVCB were involved in data analysis and interpretation. WBvH, ASJMtR, AvM, VSP, PAD, ZL, VC, PGS, and CVCB contributed reagents/materials/analysis tools. JSAvdG and AvdP drafted the manuscript. All authors critically reviewed the manuscript. All authors gave final approval of version for publication.

## Study approval

Experiments concerning human cardiac tissue were approved by the local medical ethics board, and written informed consent was provided by all patients for myocardial biopsies, UCC-UNRAVEL #12-387.^38^

## Graphical Abstract

PLN R14del cardiomyocytes demonstrate an aberrant response to mechanical stress as exemplified by 1) disrupted nuclear morphology coupled to altered nuclear calcium handling, 2) nuclear-damage induced apoptosis and strain avoidance response when exposed to cyclical stretch, and 3) disrupted mechanotransduction and cytoskeletal organisation when exposed to increased substrate stiffness, as compared to wild-type (WT) cardiomyocytes. These mechanical maladaptations of PLN R14del cardiomyocytes may contribute to the disease heterogeneity observed in PLN R14del cardiomyopathy.

## Abbreviations

ACM: Arrhythmogenic Cardiomyopathy
BSA: Bovine Serum Albumin
CTD90: Calcium Transient Duration (90%)
DCM: Dilated Cardiomyopathy
DMSO: Dimethyl Sulfoxide
GAPDH: Glyceraldehyde 3-Phosphate Dehydrogenase
hiPSC-CM: Human Induced Pluripotent Stem Cell-Derived Cardiomyocyte
HRP: Horseradish Peroxidase
HTx: Heart Transplantation
LINC: Linker of Nucleoskeleton and Cytoskeleton Complex
LMS: Living Myocardial Slices
LVAD: Left Ventricular Assist Device
OL: Overload
PBS: Phosphate-Buffered Saline
PCM-1: Pericentriolar Material 1 PFA; Paraformaldehyde
PKA: Protein Kinase A
PL: Physiological Loading
PLN: Phospholamban
PML-NBs: Promyelocytic Leukemia Nuclear Bodies
ROI: Region Of Interest
SERCA: Sarcoplasmic Reticulum Calcium-ATPase
SR: Sarcoplasmic Reticulum
TEM: Transmission Electron Microscopy
UPR: Unfolded Protein Response
WT: Wildtype
YAP: Yes-Associated Protein

