## Supplementary Data for "Increased Nuclear Susceptibility to Mechanical Stress in Phospholamban R14del Cardiomyopathy: A New Perspective on Disease Heterogeneity"

**Supplementary Figures**

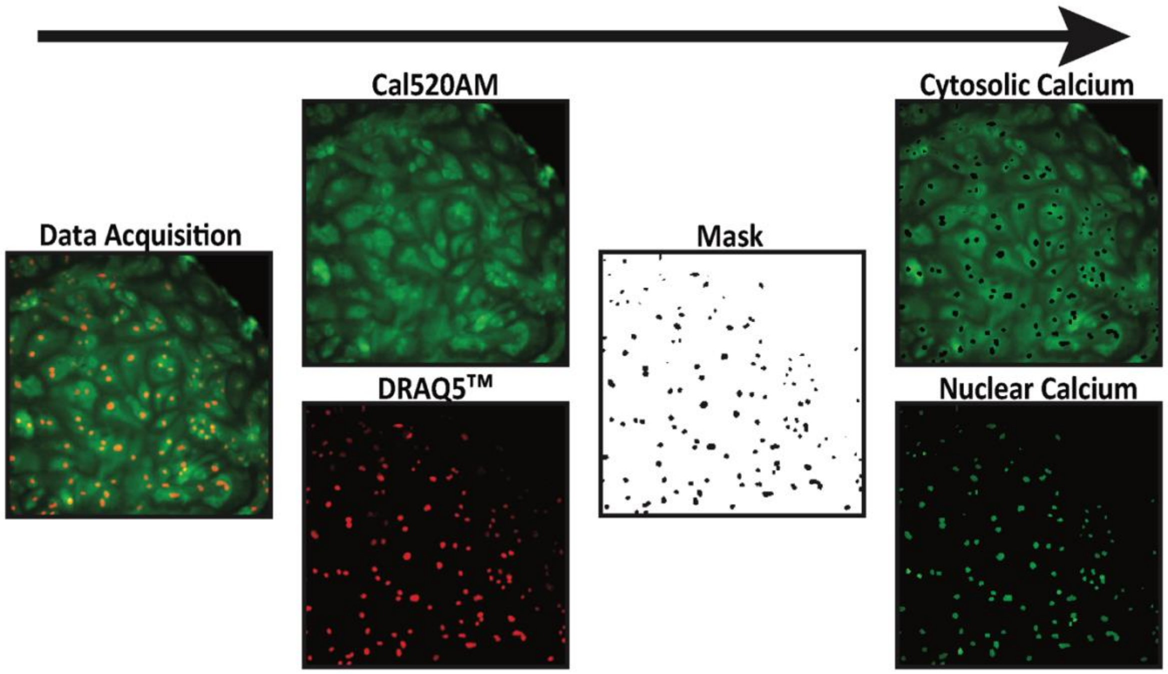


**Figure S1. Nuclear and cytosolic $Ca^{2+}$ signal segmentation workflow.** hiPSC-CMs were loaded with Cal-520AM (green) and DRAQ5™ (red) to simultaneously capture $Ca^{2+}$ dynamics and nuclear localization. Channels were separated in Fiji, and a binary nuclear mask was generated from the DRAQ5™ channel and applied to the Cal-520AM channel to isolate nuclear and cytosolic $Ca^{2+}$ signals from the same recording.

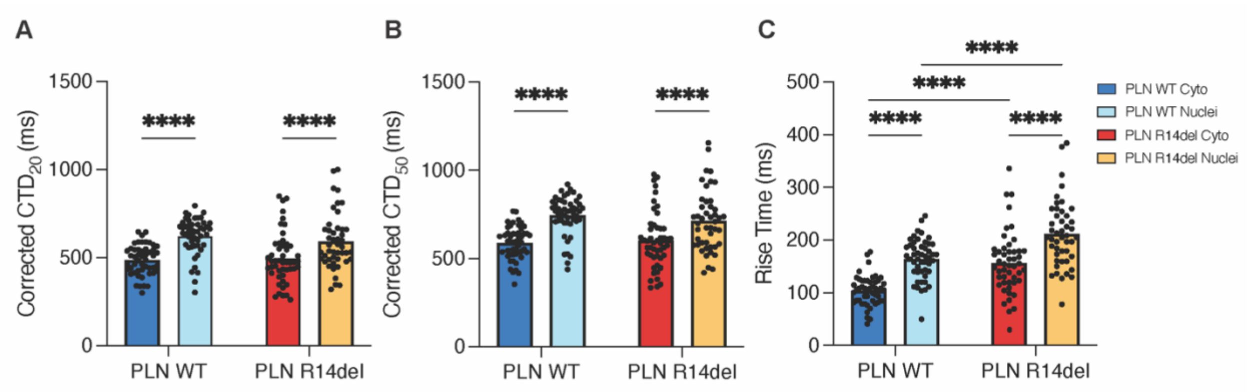


**Figure S2: $Ca^{2+}$ transient kinetics in PLN WT and R14del hiPSC-CMs.** Nuclear CTD20 (A) and CTD50 (B) are significantly prolonged compared to the cytosolic counterpart in both genotypes. C) Rise time is significantly prolonged in PLN R14del compared to PLN WT in both compartments, and significantly slower in the nuclear compared to the cytosolic compartment in both genotypes. (A-C: PLN WT n=50; PLN R14del n=50)

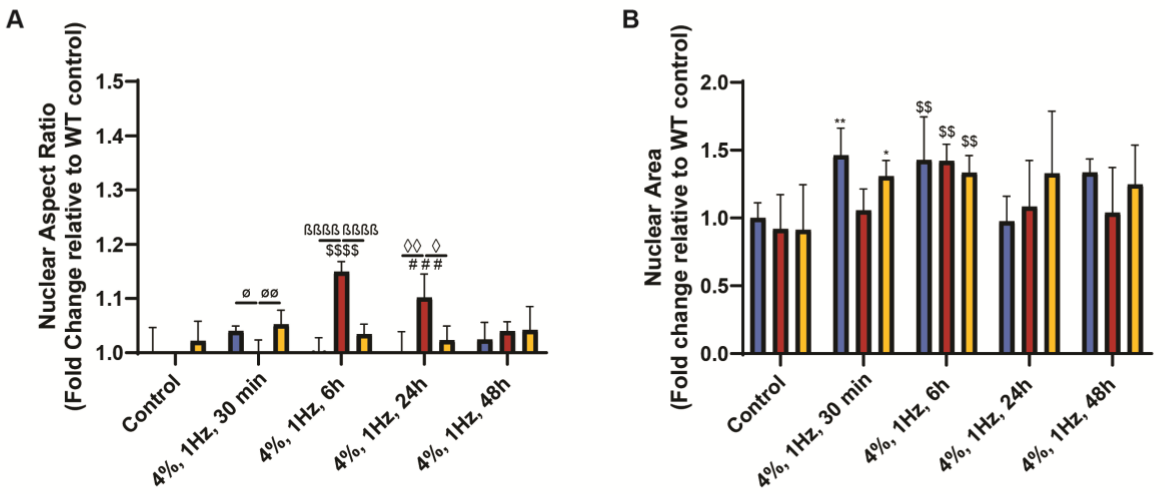


**Figure S3: Nuclear remodelling in PLN R14del hiPSC-CMs under mechanical load.** Nuclear morphology parameters, including nuclear aspect ratio (A) and nuclear area (B), were quantified as fold change relative to WT control. PLN R14del hiPSC-CMs showed an increase in nuclear aspect ratio at 6 and 24 hours of mechanical loading compared to PLN WT and PLN corr. nuclear area increased in PLN WT and PLN corr hiPSC-CMs already at 30 minutes, while all cell lines showed a significant increase at 6 hours of mechanical loading (A and B: PLN WT n=5; PLN R14del n=5; PLN corr n=5). Statistical significance is indicated by ø (30 min), β (6 h), ◊ (24 h), Δ (48 h), * (Control vs. 30 min), $ (Control vs. 6 h), and # (Control vs. 24 h).

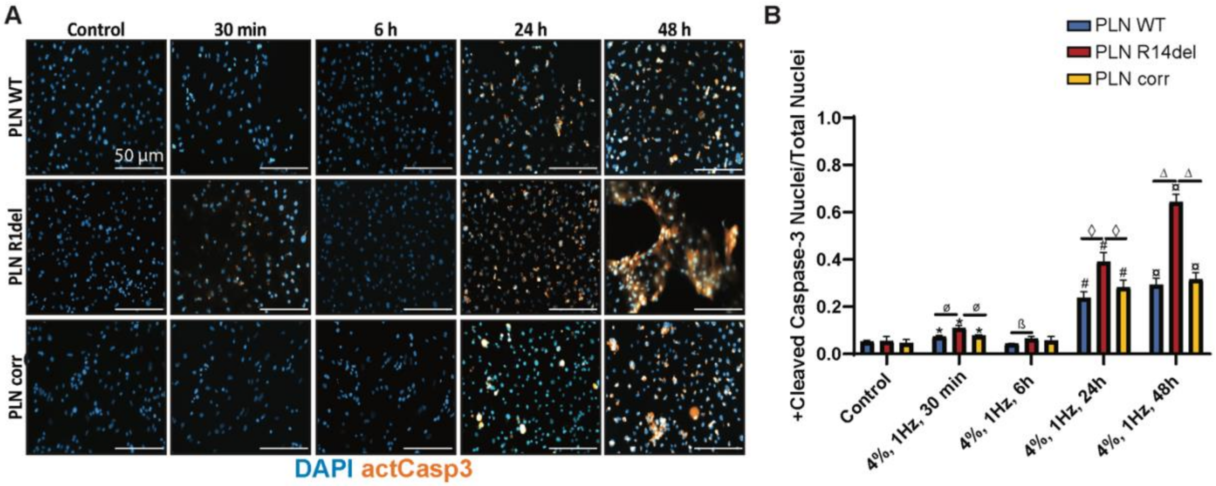


**Figure S4: PLN R14del hiPSC-CMs undergo apoptosis under mechanical load.** A) Representative immunofluorescence images of DAPI-stained nuclei (blue) and actCasp3 (orange) in PLN WT, PLN R14del, and PLN corr hiPSC-CMs in control conditions and after mechanical loading (4%, 1 Hz) for 30 min, 6 h, 24 h, and 48 h. B) Quantification of cleaved Caspase-3-positive nuclei relative to total nuclei shows a significant increase in apoptosis in PLN R14del hiPSC-CMs compared to PLN WT and PLN corr upon applying mechanical stretch. (B: PLN WT n=5; PLN R14del n=5; PLN corr n=5). Statistical significance is indicated by ø (30 min), β (6 h), ◊ (24 h), Δ (48 h), * (Control vs. 30 min), $ (Control vs. 6 h), # (Control vs. 24 h), and ¤ (Control vs. 48 h).

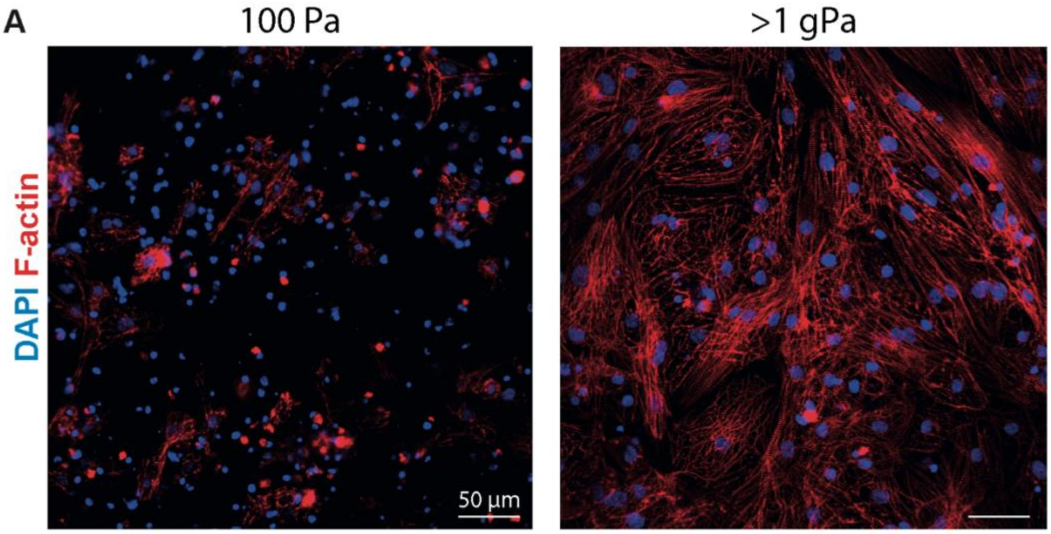


**Figure S5: Enhanced cytoskeletal organisation in cardiomyocytes with increased microenvironment stiffness.** A) Representative immunofluorescence images of DAPI-stained nuclei (blue) and F-actin (red) in hiPSC-CMs cultured on soft substrates (100 Pa) versus stiff substrates (>1 gPa). Increased substrate stiffness induces a more maturely organized actin cytoskeletal structure in hiPSC-CMs, highlighting the impact of microenvironment stiffness on cytoskeletal remodelling.

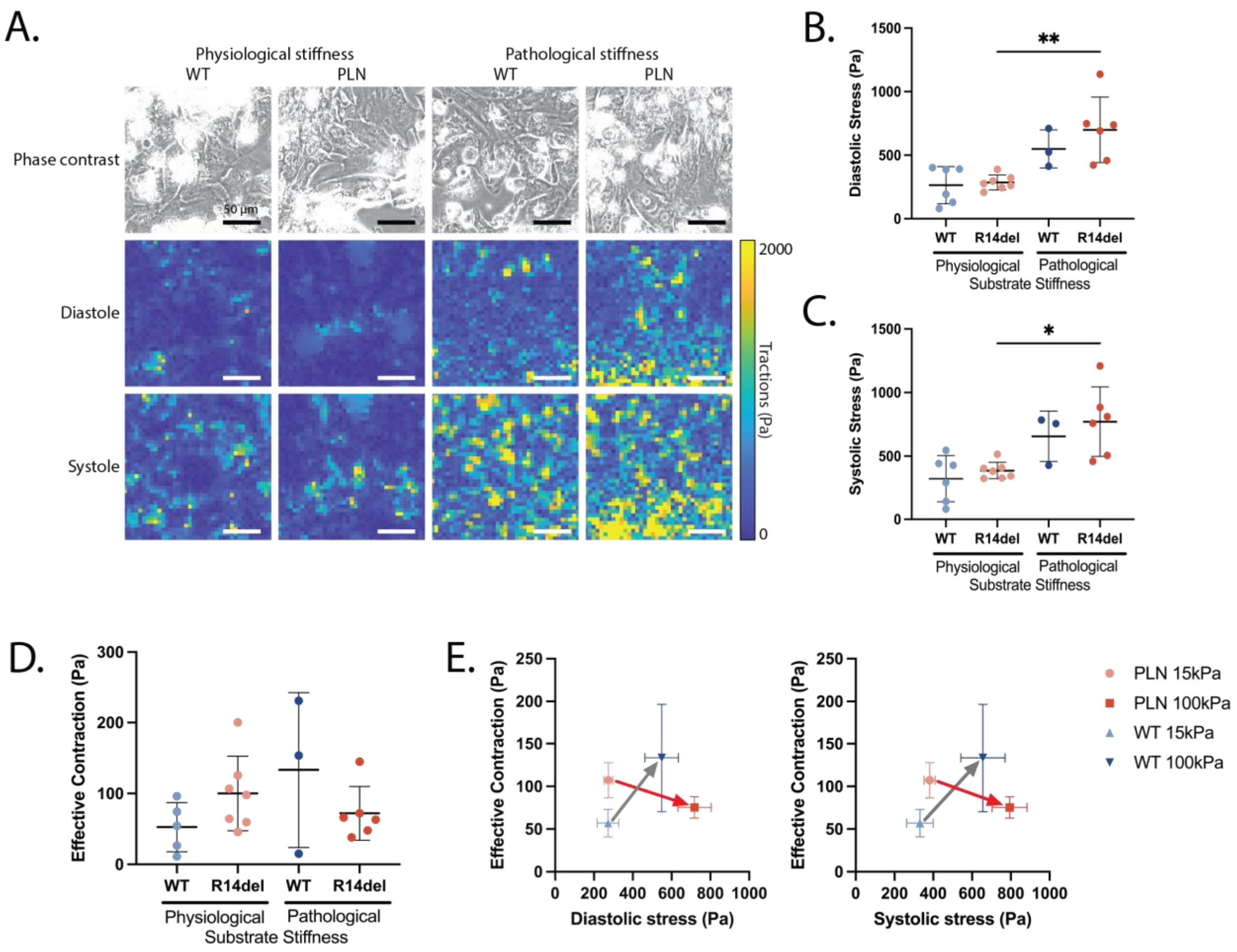


**Figure S6. PLN R14del cardiomyocytes generate elevated mechanical stress but fail to increase effective contraction on pathological substrate stiffness**. (A-C) Quantification of mechanical stress generation in PLN WT and PLN R14del hiPSC-CMs cultured on substrates with physiological and pathological stiffness. (A) Representative phase contrast images and heatmaps of traction magnitude fialds of WT and PLN cardiomyocyte monolayers on substrates of physiological and pathological stiffness. (B) Quantified diastolic mechanical stress. (C) Quantified systolic mechanical stress (WT physiological stiffness N=6, PLN R14del physiological stiffness N=7, WT pathological stiffness N=3, PLN R14del pathological stiffness N=6). Effective contraction of PLN WT and PLN R14del hiPSC-CMs on physiological and pathological stiffness substrates (WT physiological stiffness N=5, PLN R14del physiological stiffness N=7, WT pathological stiffness N=3, PLN R14del pathological stiffness N=6). (E) Relationship between mechanical stress generation and effective contraction (WT physiological stiffness N=6, PLN R14del physiological stiffness N=7, WT pathological stiffness N=3, PLN R14del pathological stiffness N=6). Data are presented as mean ± SD. Statistical significance was determined using two-way ANOVA, $^{**}P < 0.01$.

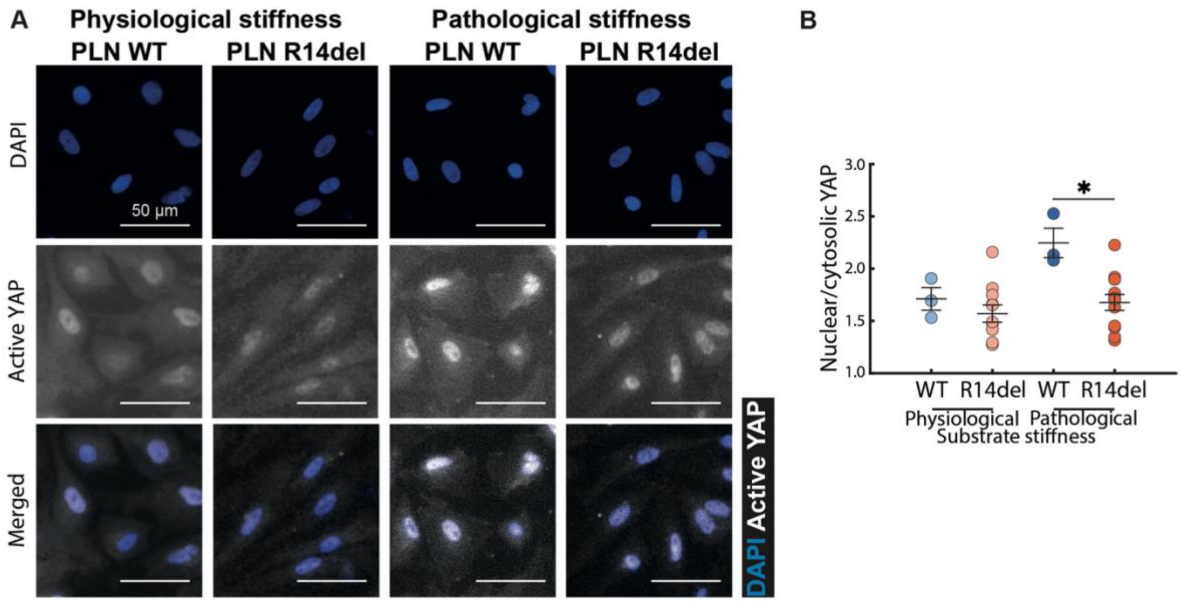


**Figure S7: Impaired YAP nuclear translocation in PLN R14del hiPSC-CMs under pathological stiffness.** A) Representative immunofluorescence images of DAPI-stained nuclei (blue) and active YAP (white) under physiological and pathological stiffness in PLN WT and R14del hiPSC-CMs. (B) Quantification of the nuclear-to-cytosolic YAP ratio reveals the inability of PLN R14del hiPSC-CMs to relocate YAP to the nucleus under pathological stiffness (15 kPa PLN WT n=3; 65-85 kPa PLN WT n=3; 15 kPa PLN R14del n=11, 65-85 kPa PLN R14del n=12).

### Supplemental Methods

#### hiPSC-CMs culture

hiPSC-CMs harbouring the PLN R14del pathogenic variant were previously described and are derived from a 52-year-old male (D4iR14del) likewise its isogenic control line, referred to as PLN corr.[1] Healthy control hiPSC line, referred to as PLN WT, has been previously registered in the European Bank for iPSCs (EBiSC, https://ebisc.org/) and was derived from a 44-year-old female UKKi032-C (NP0141-31B).[2] The maintenance of hiPSCs, their differentiation into hiPSC-CMs, and subsequent purification were performed as previously described.[3] Briefly, the cells were grown on growth-factor-reduced Matrigel (Corning) in Essential 8™ medium (Gibco), with daily medium changes. Cells were passaged every 3-4 days using 0.5 mM EDTA (Gibco). Differentiation into hiPSC-CMs was initiated by replacing the medium with 3 mL $B27^-$ medium (RPMI 1640 L-Glutamine (Gibco), 2% B27®-insulin (Gibco), and 1% Penicillin-Streptomycin (Gibco)) supplemented with 6-8 µM CHIR99021 (Selleck Chemicals) for 24 hours. On day 2, 2 mL of $B27^-$ medium was added, followed by 1 mL on day 3. On day 4, the medium was replaced with $B27^-$ medium supplemented with 2 µM Wnt-C59 (Tocris Bioscience). On day 6, the medium was changed back to $B27^-$ medium. From day 7 onwards, the cells were cultured in $B27^+$ medium (RPMI 1640 L-Glutamine, 1% B27®+insulin (Gibco), and 1% Penicillin-Streptomycin). Starting on day 9, purification was carried out, using purification medium (RPMI 1640 L-Glutamine without glucose (Gibco), 2% B27®+insulin, and 1% Penicillin-Streptomycin). On day 11, hiPSC-CMs were maintained in $B27^+$ medium, which was replaced for expansion on day 13 using a protocol involving Wnt activation by GSK-3β inhibition with 2.0 µM CHIR99021 and the inhibition of cell-cell contact.[4, 5] Subsequently, the hiPSC-CMs were cultured in $B27^+$ medium for at least 48 hours and then underwent maturation using the previously described metabolic maturation protocol for a minimum of 27 days.[6] Throughout the process, cultures were routinely screened for Mycoplasma contamination using the MycoAlert Kit (Lonza).

#### Nuclei isolation from cells and cardiac tissue

To evaluate the presence and localisation of PLN in the nucleus, nuclei were isolated from hiPSC-CMs and HeLa cells, as well as from the left ventricular tissue of PLN R14del and DCM patients. HeLa cells were used as a negative control for PLN expression. For cardiac tissue, small pieces (<1 $mm^3$) were cut and transferred to a nuclear buffer containing 0.5 mM $CaCl_2$, 0.5 mM EGTA, 2 mM EDTA, 1 mM DTT, 3 mM $MgCl_2$, and 10 mM Tris at pH 8.0. The tissue was minced using a Turrax probe homogenizer (IKA T10 basic). For cell samples, cells were detached and washed with calcium and magnesium-free PBS. Total lysate fraction (TL) was obtained by lysing cells directly in the LDS sample buffer (Thermo Fisher Scientific). Both tissue and cell samples were subsequently lysed in nuclear buffer supplemented with TritonX-100 to a final concentration of 0.2% and incubated at 4 °C for 10 minutes. Tissue lysates were sequentially passed through a 100 µm, a 70 µm, and a 40 µm cell strainer. Samples were centrifuged at 1000 x g at 4 °C for 10 minutes to separate the cytoplasmic fraction (Cyto) from the nuclear pellet. The nuclear pellet was washed twice with nuclear buffer and then pelleted by centrifugation at 1000 x g at 4 °C for 10 minutes. The purified nuclei, as well as the cytoplasmic fractions, were subsequently lysed in the LDS sample buffer.

**Western blot analysis**

Isolated proteins were analysed using Western blotting. Samples were heated at 95 °C for 10 minutes in NuPAGE reducing agent and sample buffer (Thermo Fisher Scientific). The reduced samples underwent tip sonication and were separated on a 4-12% Bis-Tris polyacrylamide gel (Thermo Fisher Scientific), alongside the PageRuler Plus Prestained Protein Ladder (ThermoFisher Scientific). The samples were transferred from the gel to a PVDF iBlot membrane (Invitrogen, iBlot Transfer Stack, PVDF) with the iBlot 2 apparatus (Life Technologies). The membrane was blocked for 1 hour with Intercept Blocking Buffer (TBS, LI-COR Biosciences). Primary antibodies, goat anti-PLN (Origene, TA311243, 1:1000, 0.5 µg/mL), rabbit anti-glyceraldehyde 3-phosphate dehydrogenase (GAPDH) (Cell Signaling, 14C10, 1:1000, 42 ng/mL), mouse anti-sarcalumenin (ThermoFisher Scientific, MA3-932, 1:1000, 1 µg/mL) and mouse anti-histone 3 (Active Motif, MABI 0301, 1: 1000), were incubated overnight at 4 °C. Secondary antibodies, Horseradish peroxidase (HRP)-conjugated rabbit anti-goat antibody (DAKO, P0449, 1:2000, 0.25 µg/mL), HRP-conjugated rabbit anti-mouse antibody (DAKO, P0260, 1:2000, 0.65 µg/mL), HRP-conjugated goat anti-rabbit antibody (DAKO, P0448, 1:2000, 0.13 µg/mL), were incubated for 1 hour at room temperature. The primary and secondary antibodies were diluted in Intercept Blocking Buffer (TBS). Proteins were detected with a chemiluminescent peroxidase substrate (Sigma-Aldrich) and a ChemiDoc XRS+ system (Bio-Rad) with Image LabTM software.

**Nuclear calcium imaging**

Matured hiPSC-CMs were incubated with 2.5 mM DRAQ5™ (ThermoFisher Scientific), 1.25 mM Cal-520AM (Abcam, ab171868), and 0.5% Pluronic F-127 (Sigma-Aldrich) in 50% Fluorobright DMEM (ThermoFisher Scientific) for 30 minutes at 37 °C. After incubation, the plate was transferred to a 37 °C preheated Thunder DMi8 (Leica) microscope, and a 30-second recording of Cal-520AM fluorescence was acquired, followed by a DRAQ5™ fluorescence image to serve as a nucleus-selective mask, as illustrated in Figure S1. Fiji 2.14.0/1.54f software was used to separate nuclear and cytosolic calcium transients. Peaks, a custom-written MATLAB script (DOI:10.17605/OSF.IO/86UFE), was used to analyze the calcium transients. 20%, 50% and 90% of the calcium transient duration ($CTD_{20}$, $CTD_{50}$, $CTD_{90}$ respectively), for both nuclear and cytosolic calcium, was corrected for spontaneously beating using a modified Fredericia's correction: corrected $CTD_x = CTD_x/(\sqrt[3]{}(60/\text{beats per minute})$ as described before.[7, 8] Rise time was additionally calculated for both compartments.

**Mechanical stretching of hiPSC-CMs**

One week before initiating the experiments, hiPSC-CMs were detached using 1x TrypLE (Gibco) and plated using replating medium on Matrigel (Corning, 356231) coated 6-well BioFlex® Flexible-bottom culture plates (Flexcell Inc., Burlington, USA). After 24 hours, the medium was replaced with $B27^+$ medium for an additional 48 hours. Subsequently, the cells were maintained in maturation medium for experimental procedures. The FX-5000™ Tension System (Flexcell Inc., Burlington, USA) was used to provide controlled uniaxial cyclic strain to the cardiomyocytes cultured on the flexible membranes using a regulated vacuum pressure that deforms the membrane over a rectangular post underneath the membrane.[9] Cardiomyocyte cultures were subjected to 4% cyclic strain at 1 Hz for 30 minutes, 6 hours, 24 hours, or 48 hours. Time-matched unstrained cardiomyocytes served as controls.

**Histological analysis of hiPSC-CMs**

For immunofluorescence staining following both mechanical stretch and substrate stiffness experiments, cells were washed three times with PBS before being fixed in 3.7% (v/v) formaldehyde in PBS (Sigma-Aldrich) for 10 minutes at room temperature. Cells were permeabilized with 0.5% Triton-X-100 (Merck) in PBS for 10 minutes and blocked with 5% bovine serum in PBS for 30 minutes at room temperature. After blocking, cells were incubated with primary antibodies diluted in washing buffer for 1.5 hours at room temperature: rabbit anti-cleaved Caspase-3 (9884S, Cell Signaling Technology, 1:500, 2.6 μg/mL), mouse anti-promyelocytic leukaemia nuclear bodies (PML-NB) (SC-966, Santa Cruz, 1:200, 1 μg/mL), rabbit anti-Desmin (D8281, Sigma Aldrich, 1:100, 0.82 μg/mL), rabbit anti-Nesprin-1 (PA5-115640, Thermo Fischer, 1:100, 10 μg/mL), TUNEL (A23210, Invitrogen, per manufacturer's instructions), and mouse anti-Yes-associated protein (YAP) (SC-101199, Santa Cruz, 1:200, 0.5 μg/mL). After the primary incubation, cells were subjected to three PBS washes for 5 minutes, followed by a 1.5-hour incubation with secondary antibodies diluted in the washing buffer: goat anti-mouse Alexa488 (A21131, Molecular Probes, 1:300, 6.67 μg/mL), goat anti-mouse Alexa488 (A21121, Molecular Probes, 1:500, 4 μg/mL), donkey anti-rabbit Alexa555 (A31572, Invitrogen, 1:1000, 2 μg/mL), and Phalloidin 647N (D65906, Sigma Aldrich, 1:200). The cell nuclei were then stained with DAPI (D9542, Sigma Aldrich, 1:500) for 5 minutes, following three washing steps with the washing buffer. Mechanical stretch samples were then mounted onto a microscope glass slide using a Mowiol-based mounting medium, while substrate stiffness samples were covered in PBS and stored at 4 °C until image acquisition.

Image acquisition was performed with the Zeiss Axio Observer 7 equipped with an LD Plan-Neofluar 20x/0.4 Korr M27 objective. To obtain Z-stack images, used for quantifying the nuclear and actin height, the Leica SP5x confocal microscope, equipped with an HCX PL APO CS 20.0x0.70 objective, was used. The Z-stacks were approximately 20 μm thick (50-60 slices).

**Image processing**

For all image quantifications of hiPSC-CMs, we used the StarDist tool and skimage in Python.[10] StarDist detected and segmented nuclei in the images. Using skimage.measure, we calculated morphological properties, including nuclear and cellular area, and the aspect ratio. The aspect ratio was defined as the ratio between the major and minor axis lengths, providing a measure of how elongated cells and nuclei are. Orientation was determined for both the nucleus and the cytoskeleton as the angle between the major axis of the fitted ellipse and the horizontal axis, normalized to a range of 0 to 180 degrees.

To assess nuclear susceptibility, we determined the amount of nuclear injury marker PML-NB within the nuclei. StarDist segmented nuclei from DAPI-stained images, and PML-NBs were identified. Only PML-NBs overlapping with nuclei and within a size range of 10 to 150 pixels were included. Nuclear susceptibility was determined by counting the total number of PML-NBs within each nucleus and calculating the ratio of the total PML-NBs to the total nuclei. Similarly, the amount of cleaved Caspase-3 and TUNEL-positive cells per image was counted using StarDist in Python. Blobs and nuclei were segmented with StarDist, and only blobs that overlapped with the nucleus or were within a maximum distance threshold were counted as cleaved Caspase-3 or TUNEL-positive cells. To determine the total cytoplasmic levels of Desmin and Nesprin-1, we used a CellProfiler workflow to measure the total intensity of their respective staining. To analyse perinuclear Desmin and Nesprin-1, staining intensity was quantified within a region of interest (ROI) defined as an 8-pixel-wide band surrounding the nuclear boundary.

The number of F-actin fibres was quantified in MATLAB using the segmentfibers script (https://github.com/tkphung/SegmentFibers). In brief, pixels that belonged to an F-actin fibre were identified using the Frangi Vesselness.[11] The number of F-actin fibres was then quantified as the ratio between pixels containing fibre and the total pixel count, which was normalized by the number of nuclei in the image.

All quantifications of F-actin and nucleus height were performed in MATLAB using a custom script. Orthogonal XZ projections of nuclei and F-actin were binarized using Otsu's method. In the resulting binary images, the vertical distance between the highest and lowest pixel values was calculated. For nuclear height, a Gaussian function was plotted (with 1, 2, or 3 terms, depending on the number of nuclei present in the image) and fitted to the nuclear height profile. The peak values were used to determine the individual nuclear heights. To determine total cell height, the same vertical distance calculation was applied to the F-actin channel. For the actin fibre height specifically, the average of the measured distances across the image was used as quantification.

**Preparing Polyacrylamide (PAA) substrates for stiffness sensing by hiPSC-CMs**

To prepare PAA substrates of distinct stiffness, a 40% (w/v) acrylamide solution (Bio-Rad) and 2% bis-acrylamide solution (Bio-Rad) were mixed with phosphate-buffered saline (PBS) (Sigma-Aldrich) in different ratios, as indicated in Table 1. To facilitate the crosslinking of acrylamide and bis-acrylamide, these solutions were supplemented with 0.5% of 10% ammonium persulfate (Bio-Rad) and 0.05% of N,N,N,N'-tetramethylethylenediamine (Sigma-Aldrich). Finally, to determine substrate deformation for traction force microscopy, 1% of FluoSpheres carboxylate modified microspheres (0.2 mm, dark red; ThermoFisher Scientific) was added. Concurrently, glass-bottom 12-well plates (No. 0 thickness, MatTek) were treated with a 14:1:1 solution of absolute ethanol

(VWR), acetic acid (Merck), and PlusOne Bind-Silane (VWR) for 1 hour at room temperature, to enable the adhesion of the PAA substrates to the glass bottoms. Droplets of 11.5 µl PAA solution were pipetted into the 12-well plates and subsequently covered with 13 mm-diameter coverslips (No. 1 thickness, VWR). Hydrogel substrates were polymerized during a 1-hour incubation step at room temperature, after which PBS was added to each well. The glass coverslips were carefully removed with a needle, and the PAA substrates were washed twice with PBS prior to collagen coating and cell seeding.

Mechanical characterisation of the substrates was performed using the Piuma Nanoindenter (Optics 11) equipped with probes of 0.34 N/m spring constant and a spherical tip radius of 24 µm. For each substrate, the stiffness was assessed at four locations in the mid-section of the gel, forming a square of 2x2 mm. Each measurement included a 7-µm indentation executed over 2 seconds, which was subsequently maintained for 2 seconds before being retracted in 2 seconds. The bulk substrate stiffness was calculated using a Hertzian contact model, fitted to the load-displacement curve. The PAA substrates were assumed to be quasi-incompressible and thus a Poisson's ratio of 0.49 was employed in the model. The characterisation was conducted on substrates without a collagen coating, as the effect of this coating on the bulk stiffness of the substrate is negligible (data not shown)[12].

A collagen IV coating was applied to the PAA substrates to facilitate cell attachment. To this end, PAA substrates were treated with Sulfo-SANPAH (ThermoFisher Scientific) for 5 minutes, while exposed to UV light (365 nm wavelength, XX-15, UVP), enabling collagen adherence to PAA. After three washing steps with sterile PBS to remove the Sulfo-SANPAH solution, the substrate was coated with human collagen type IV (0.1 mg/ml solution, Sigma-Aldrich). Collagen droplets were incubated overnight at 4 °C and washed off with sterile PBS before cell seeding.

**Seeding and culturing hiPSC-derived cardiomyocytes on PAA substrates**

PAA substrates were covered with a replating medium (RPMI 1640 L-Glutamine supplemented with 10 µM ROCK inhibitor (Sigma-Aldrich), 10% KOSR, 1% B27®+insulin, and 1% Penicillin/streptomycin) droplet using a pipette and incubated at 37 °C for 1 hour. During this incubation step, PLN R14del and PLN WT cardiomyocytes were harvested using 1x TrypLE (Gibco) and seeded in droplets of maturation medium, each containing 80.000 cells, onto the PAA substrates. Cells were allowed to attach to the substrate during a 1-hour incubation step at 37°C. Next, 1 ml of replating medium was gently added to each well. Cells were cultured on substrates for 1 day before replacing the replating medium with 2 ml of maturation medium.

**Time-lapse microscopy**

Time-lapse microscopy was performed using an inverted microscope (Eclipse Ti2, Nikon), equipped with a 20x/0,75 CFI Plan Apo λ DM Ph2 objective (Nikon). This microscope featured an incubation chamber, with temperature set at 37 °C, CO2 level of 5%, and 95% humidity. High speed imaging bursts of ~4,3 seconds were acquired in series of the cells, of the dark red-fluorescent microspheres in the PAA substrate, and optionally of the green-fluorescent calcium dye. Imaging of cells was performed with phase contrast microscopy, whereas the microspheres and, if applicable, the calcium dye were imaged with epifluorescence microscopy. The imaging bursts were captured in distinct regions of interest (ROI) every 30 minutes over a total period of 2 hours. Up to 60 ROIs, constituting replicates, spread over 12 substrates were imaged using a motorized stage, NIS software, and perfect focus system. Following a time-lapse experiment, cells were dissociated using a 10% SDS (Merck) solution, and reference images of the fluorescent microbeads in the undeformed substrates were acquired. These reference images enabled the computation of cell-induced deformation of the substrate for traction force microscopy analyses. Distinct time-lapses on different days with distinct batches of PAA substrates and cell passages were considered repeats.

**Traction force microscopy**

Post processing and analyses of traction force microscopy-related images and data were performed in MATLAB (MathWorks) with custom codes. To obtain the displacement fields of the PAA substrates in each ROI, particle image velocimetry using PIVlab[13] was performed (32-pixel resolution, 0,5 overlap) on the fluorescent microbead images from each burst at each time-point of the time-lapse and the corresponding reference image. Displacement fields were converted into traction fields using a custom script that applies a Fourier transform algorithm assuming an elastic hydrogel with finite thickness.[14] To purge edge effects, traction fields were cropped by 5% on each side. Cell masks were acquired using the first phase contrast image of the cells in each burst, which were used to isolate the traction values in the field corresponding with cells. To assess cardiomyocyte function, the median traction magnitude within the cell mask was computed for each frame of a burst. The baseline mechanical stress during diastole was computed by applying a 20th-order median filter to the median traction magnitude during a burst. The peaks of maximum stress of cardiomyocytes during systole were identified by applying MATLAB's findpeaks function on the median traction magnitude during each burst. Peaks were included based on three criteria; (1) their magnitude exceeded the baseline mechanical stress by 5% plus two times the standard deviation of the baseline, (2) they lasted more than 0.04 seconds, (3) neighboring peaks were more than 0.4 seconds apart. The effective contraction of cardiomyocytes was determined as the difference in average systole peak magnitude and the diastolic mechanical stress within a burst. Per ROI, the diastolic mechanical stress, systolic mechanical stress, and effective contraction per burst were averaged per biological repeat.

**Generating living myocardial slices and mechanical overloading**

LMS were prepared as described before.[15] In brief, a fresh myocardial biopsy was obtained from the left ventricle of a 52-year-old male end-stage heart failure patient carrying the PLN R14del variant at the time of heart explantation. The myocardial biopsy was transported in an ice-cold modified Tyrode's solution (30 mM BDM, 0.9 mM $CaCl_2$, 10 mM glucose, 10 mM HEPES, 9 mM KCl, 1 mM $MgCl_2$, and 140 mM NaCl at a pH of 7.4), mounted with the epicardial side facing down and sectioned into 300 μm-thick LMS using a high-precision vibratome (7000 smz-2,

Campden Instruments Ltd, Gillingham, UK). The vibratome was set to a z-axis error of <1.0 μm, a frequency of 80 Hz, 2 mm amplitude, and an advance speed of 0.03 mm/s. The slices were trimmed and attached perpendicular to the myofibril direction using 0.1 mm-thick polyester copier clear triangles and histoacryl glue (B. Braun, Melsungen AG, Germany). LMS were placed in biomimetic culture chambers (InVitroSys, München, Germany), where they were continuously electrically paced (3 ms biphasic 50 mA pulses, at 0.5 Hz), stretched to a physiological stretch of 19.6%[16], and rocked at 60 rpm. LMS were cultured in medium 199 supplemented with 1% ITS, 0.2% Primocin, 2.15 nM Triiodothyronine (T3), 100 nM Dexamethasone, 4 nM Adrenaline, 4 nM Noradrenaline, 50 μM 2-Mercaptoethanol, and 20 μg/mL Ascorbic Acid (all from Sigma-Aldrich). Media was changed every 48 to 72 hours, and conditioned media was collected for further analysis. Experiments commenced after a 1-week adaptation period. LMS were either maintained at a physiological stretch of 19.6% (PL) or subjected to mechanical overloading (OL) by stretching to 29% of the original slice length, as previously described[17]. Subsequently, LMS were fixed in 4% paraformaldehyde (PFA) and embedded in paraffin.

**Histological analysis of cardiac tissues**

To assess the effect of mechanical loading on nuclear morphology in patients, cardiomyocyte nuclei were analysed in paraffin-embedded tissue sections derived from fixed LMS and cardiac biopsies. Myocardial biopsies were obtained from three patients carrying the PLN R14del variant, three patients with dilated cardiomyopathy (DCM) of suspected but unidentified genetic origin despite comprehensive genetic screening, and three non-failing donor hearts that were not used for transplantation due to technical reasons, serving as healthy controls. For each patient, paired myocardial samples were analysed at two points: apical biopsy collected at the time of LVAD implantation (mechanical loaded state), and a left ventricular free wall sample collected at heart transplantation (HTx) (after mechanical unloading). This design enabled direct within-patient comparison of nuclear morphology under differing mechanical conditions.

Paraffin-embedded samples of fixed LMS and cardiac tissues were sectioned at a thickness of 4 μm for histological analysis. First, sections were deparaffinized and subjected to antigen retrieval using a citrate buffer at pH 6.0. After cooling, sections were blocked using 5% BSA for 1 hour at room temperature and incubated overnight at 4 °C with the primary antibodies in 1% BSA in PBS: mouse anti-PLN (2D12, Thermo Fisher Scientific, 1:100, 10 μg/mL) and rabbit anti-PCM-1 (HPA023370, Sigma-Aldrich, 1:500, 0.4 μg/mL). Note: the PLN 2D12 antibody epitope encompasses residue 14 of PLN, and quantitative comparison between PLN WT and PLN R14del protein expression is therefore not possible. After washing with 0.5% Tween-20, sections were incubated for 1 hour at room temperature with DAPI (Thermo Fisher Scientific, 1:2000) and secondary antibodies: donkey anti-rabbit Alexa Fluor 488 (A21206, Thermo Fisher Scientific, 1:400, 5 μg/mL), donkey anti-rabbit Alexa Fluor 555 (A31572, Thermo Fisher Scientific, 1:400, 5 μg/mL), and donkey anti-mouse Alexa Fluor 555 (A31570, Thermo Fisher Scientific, 1:400, 5 μg/mL). Finally, the stained sections were washed and mounted with a coverslip using Mowiol 4-88 (Sigma-Aldrich). Image acquisition was conducted using a NanoZoomer S360 Digital slide scanner (Hamamatsu Photonics K.K, Hamamatsu, Japan). Image assembly, merging, and contrast/brightness adjustments were performed in Fiji 2.14.0/1.54f software.

**Ultrastructural analysis of nuclei in tissue using transmission electron microscopy (TEM)**
Fresh cardiac biopsies acquired at LVAD implantation and heart transplantation were fixed in modified Karnofsky fixative (2% paraformaldehyde, 0.1% glutaraldehyde in phosphate buffer) overnight at 4 °C and post-fixed for 2 hours with 1% osmium tetroxide ($OsO_4$) in 0.1 M phosphate buffer at a pH of 7.4. After fixation, samples were dehydrated through a graded acetone series and embedded in epoxy resin (EPON). Ultrathin sections were cut using a Leica Ultracut UCT ultramicrotome, mounted on nickel grids, and stained with uranyl acetate and lead citrate using a Leica AC20 staining system. Micrographs were obtained using either a JEOL1010 TEM (JEOL Ltd., Tokyo, Japan) at 80 kV or a Tecnai12 TEM (FEI/Thermo Fisher Scientific), both equipped with a Veleta 2k × 2k CCD camera M (EMSIS, Münster, Germany).

**Analysis of cardiomyocyte nuclei in tissue**
For histological analysis, five longitudinal sections containing cardiomyocytes were selected from transmural paraffin-embedded tissue derived from both LMS and cardiac biopsies obtained during LVAD implantation and HTx. Nuclear segmentation and analysis were performed using Imaris software (Bitplane, Oxford Instruments, Belfast, UK, v10). Segmentation of PCM-1- and DAPI-positive nuclei was achieved through machine learning algorithms, followed by blinded manual refinement of the data to ensure data accuracy and quality. For each identified nucleus, the nuclear area, circularity, and dimensions of the bounding box were measured. Nuclear eccentricity was derived from bounding box axes as a measure of nuclear deformation.
Given the nuclear envelope's critical role in maintaining nuclear integrity and mechanotransduction, evaluating its morphological regularity provides insight into nuclear stress and potential disruptions in nuclear-cytoskeletal coupling[18]. To evaluate the regularity of the nuclear envelope, the total length of the nuclear envelope was manually measured from TEM images using Fiji software (v2.14.0/1.54f). This length was normalized to the circumference of the nucleus in the absence of invaginations.

**Ultrastructural analysis of nuclei in tissue using transmission electron microscopy (TEM)**
Fresh cardiac biopsies acquired at LVAD implantation and heart transplantation were fixed in modified Karnofsky fixative (2% paraformaldehyde, 0.1% glutaraldehyde in phosphate buffer) overnight at 4 °C and post-fixed for 2 hours with 1% osmium tetroxide ($OsO_4$) in 0.1 M phosphate buffer at a pH of 7.4. After fixation, samples were dehydrated through a graded acetone series and embedded in epoxy resin (EPON). Ultrathin sections were cut using a Leica Ultracut UCT ultramicrotome, mounted on nickel grids, and stained with uranyl acetate and lead citrate using a Leica AC20 staining system. Micrographs were obtained using either a JEOL1010 TEM (JEOL Ltd., Tokyo, Japan) at 80 kV or a Tecnai12 TEM (FEI/Thermo Fisher Scientific), both equipped with a Veleta 2k × 2k CCD camera M (EMSIS, Münster, Germany).
